# SCING: Single Cell INtegrative Gene regulatory network inference elucidates robust, interpretable gene regulatory networks

**DOI:** 10.1101/2022.09.07.506959

**Authors:** Russell Littman, Ning Wang, Chao Peng, Xia Yang

## Abstract

Gene regulatory network (GRN) inference is an integral part of understanding physiology and disease. Single cell/nuclei RNAseq (scRNAseq/snRNAseq) data has been used to elucidate cell-type GRNs; however, the accuracy and speed of current scRNAseq-based GRN approaches are suboptimal. Here, we present Single Cell INtegrative Gene regulatory network inference (SCING), a gradient boosting and mutual information based approach for identifying robust GRNs from scRNAseq, snRNAseq, and spatial transcriptomics data. Performance evaluation using held-out data, Perturb-seq datasets, and the mouse cell atlas combined with the DisGeNET database demonstrates the improved accuracy and biological interpretability of SCING compared to existing methods. We applied SCING to the entire mouse single cell atlas, human Alzheimer’s disease (AD), and mouse AD spatial transcriptomics. SCING GRNs reveal unique disease subnetwork modeling capabilities, have intrinsic capacity to correct for batch effects, retrieve disease relevant genes and pathways, and are informative on spatial specificity of disease pathogenesis.

## Introduction

Understanding pathophysiology is necessary for the diagnosis and treatment of complex diseases, which involve the perturbation of hundreds or thousands of genes^1–4^. Identifying perturbed gene pathways and key drivers of complex diseases requires the elucidation of gene regulatory networks (GRN) from high dimensional omics data^5,6^. Previous approaches have been developed and applied to identify these GRNs through bulk transcriptomic data and to determine causal mechanisms of disease^7–9^. More recently, with the advent of single cell RNA sequencing (scRNAseq) and spatial transcriptomics, the contributions of numerous genes in individual cell types have been implicated in diseases across many disciplines of biology and medicine^10–12^.

GRN construction from scRNAseq data has been tackled with limited success^13–15^. Existing GRN tools utilize scRNAseq data with thousands of pre-select genes and cells, because GRNs from full transcriptomes in large scRNAseq datasets are often computationally intensive and intractable^13,15^. Additionally, benchmarking studies have shown limited accuracy of existing methods on both synthetic and real data^13^. While tools such as ppcor^16^ and PIDC^17^ use partial correlation and partial information decomposition, respectively, to identify gene coexpression modules, few methods are able to identify directed networks^14^. One tool that can identify directed networks is GRNBOOST2^18^, a gradient boosting based method that has been successful in benchmarking studies^13,15^. However, most of the regulatory edges from GRNBOOST2 point in both directions, and the resulting GRNs contain too many edges in the range of 34,000 to 47,000 edges for networks with only 3,000 input genes, making this approach impractical for datasets with thousands of cells and full transcriptomes^18^. SCENIC^19^, an extension of GRNBOOST2, prunes edges based on known transcription factor binding sites (TFBS). However, it only focuses on regulatory behavior between transcription factors (TF) and its downstream target genes^19^, thereby missing other non-TF gene regulatory mechanisms^20,21^, as well as regulatory patterns due to low TF expression^22^. While there exist other GRN inference approaches based on pseudotime analysis^23–25^, their performance is generally inferior to those that rely solely on the scRNAseq data^13^.

Here, we present Single Cell INtegrative gene regulatory network inference (SCING), a gradient boosting based approach to efficiently identify GRNs in full single cell transcriptomes. The robustness of SCING is achieved via i) merging and taking consensus of GRNs through bagging, and ii) further directing and pruning edges through the use of edge importance and conditional mutual information. SCING GRNs are then partitioned into modules, to compute module specific expression for each cell. These modules can be used for clustering, phenotypic association, and biological annotation through pathway enrichment. We show our approach is both efficient and robust on large scRNAseq datasets, able to predict perturbed downstream genes of high throughput perturbation experiments, and produces gene subnetworks with biologically meaningful pathway annotations. We evaluate our approach against GRNBOOST2, ppcor, and PIDC through perturbation target prediction in Perturb-seq data, goodness of fit, network characteristic metrics, and disease modeling accuracy. Furthermore, we apply SCING to the mouse single cell atlas^26^, snRNAseq^27^, and spatial transcriptomics^28^ datasets to demonstrate its versatility in datatype accomodation and its biological interpretability of high-throughput transcriptomics datasets. Our code and tutorials for running SCING are publicly available at https://github.com/XiaYangLabOrg/SCING.

## Results

### SCING method and evaluation overview

SCING leverages the power of abundant cell-level transcriptome data from scRNAseq/snRNAseq to identify potential directional regulatory patterns between genes. However, single cell transcriptomics data has technical issues, such as high sparsity and low sequencing depth^29,30^, which make the use of traditional linear or correlative approaches challenging. Additionally, these datasets often contain tens of thousands of cells each with hundreds to thousands of genes, making identifying GRNs using complex non-linear approaches on full transcriptomes difficult^13^. To address these limitations, we employ a combination of supercell, gene neighborhood based connection pruning, bagging, gradient boosting regression, and conditional mutual information approaches to identify robust regulatory relationships between genes based on single cell transcriptomics data (Figure 1a; Methods).

**Figure 1.**
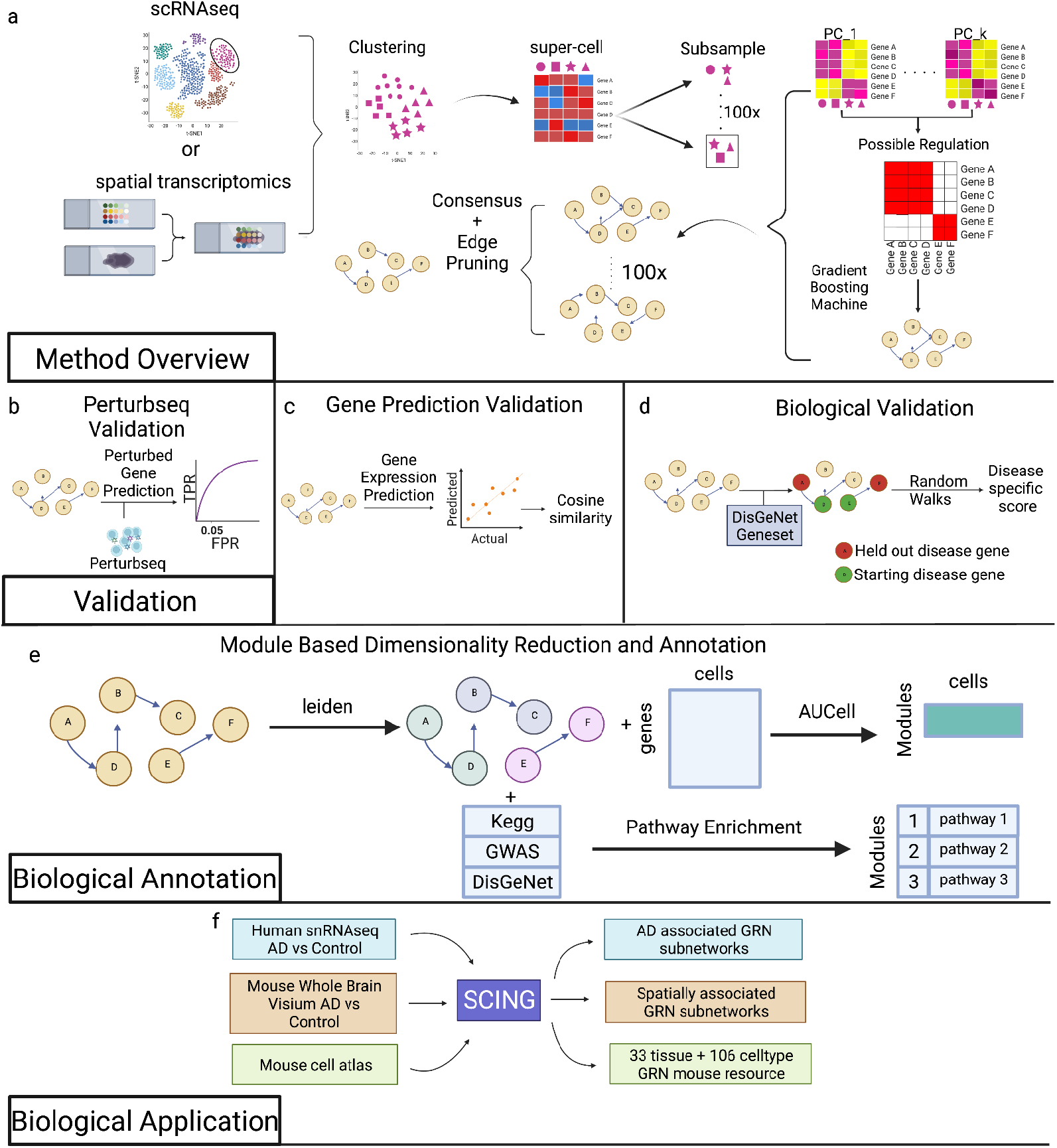
SCING overview, benchmarking, and application. SCING overview (a). First, we select a specific cell type, or use whole visium data. We then cluster the cells using the leiden graph partitioning algorithm and merge subclusters into supercells. We utilize bagging through subsamples of supercells to keep robust edges in the final GRN. For each subsample, the genes are clustered to only operate on likely regulatory edges. We then identify edges through gradient boosting regressors (GBR). We find the consensus as edges that show up in 20% of the networks. We then prune edges and cycles using conditional mutual information petrics. perturb-seq validation (b). We identified downstream perturbed genes of guides for specific genes. We then predict perturbed genes at each depth in the network from the perturbed gene. True positive rate, and false positive rate are determined at each depth in the network. We utilize AUROC and TPR at FPR 0.05 as metrics for evaluation. Gene prediction validation, to determine network overfitting (b). We split data into train and test sets and build a network on the train set. A GBR is trained for each gene based on its parents in the train data. We then predict the expression of each gene in the test set and determine the distance from the true expression through cosine similarity. Biological validation through disease subnetwork modeling (c). We utilize a random walk framework from Huang et al. to determine the increase in performance of a GRN to model disease subnetworks versus random GRNs with similar node attributes (d). We utilize the leiden graph partitioning algorithm to identify GRN subnetworks. We combine these subnetworks with the AUCell method to get module specific expression for each cell and further combine the gene modules with pathway knowledge bases to annotate modules with biological pathways (e). We apply SCING to human prefrontal cortex snRNAseq data with AD and Control patients, whole brain visium data, for AD vs WT mice at different ages, and to the mouse cell atlas in 33 tissues and 106 cell types (f).

We evaluated our approach against PIDC, a partial information decomposition approach; ppcor, a partial correlation approach; and GRNBOOST2, a gradient boosting approach. We selected these particular methods for comparison due to their overall better performance in recent benchmarking studies^13,15^ and diverse approaches. We compared these methods on the ability to predict downstream gene targets of large scale perturb-seq studies (Figure 1b), robustness of the network on training and test data (Figure 1c), metrics including the consistency of edge overlap on GRNs built on independent cells, and the ability to model disease subnetworks (Figure 1d). Furthermore, we demonstrated the utility of using SCING on the full mouse cell atlas and a human prefrontal-cortex snRNAseq dataset^27^ with AD and control patients to perform snRNAseq batch harmonization, gene module identification with biological annotation (Figure 1e), and module-trait association analysis (Figure 1f). Data from Morabito et al. has snRNAseq from 11 AD and 7 control human prefrontal cortex samples, with 61,472 nuclei across 7 cell types, which provides a high quality dataset for benchmarking. Furthermore, we applied SCING to a visium mouse dataset with AD vs control samples^28^. We show that the SCING subnetworks are versatile in data type accomodation (scRNAseq, snRNAseq, spatial transcriptomics), can resolve spatial biology, and are powerful in retrieving biologically meaningful pathways, gene connections, and disease associations.

### SCING extends network node inclusion capacity and improves computing speed

SCING builds many GRNs for each dataset and the speed of such computation is paramount to reasonable computation for a whole dataset. The use of supercells and gene covariance based potential edge pruning enables faster performance of SCING. We show that SCING improves computational speed over GRNBOOST2 and PIDC, when increasing the number of genes (SF1a, Table 1), and number of cells (SF1b, Table 2). GRNBOOST2 scales exponentially on the number of genes, while PIDC scales exponentially on the number of cells, making whole transcriptome and large dataset GRN inference difficult. Supercells in SCING ensure the network building run time does not increase as a function of cells and potential edge pruning enables linear increase in computation with respect to genes. We note that ppcor’s fast general matrix formulation improves GRN inference time compared to all other approaches, including SCING. While SCING is slower than ppcor, it performs inference on 4,000 genes in ~21 seconds for all cell types, which is reasonable to compute hundreds of GRNs for any given sample.

**Table 1.** Average run time (seconds) of GRN building methods on variable number of genes for 1,000 cells across 10 iterations.

|  |  | 1000 genes | 2000 genes | 4000 genes |
| --- | --- | --- | --- | --- |
| astrocytes | SCING | 10.8 | 14.2 | 21.4 |
|  | ppcor | 4.3 | 4.1 | 4.1 |
|  | PIDC | 74.3 | 77.2 | 85.1 |
|  | GRNBOOST2 | 35.0 | 98.2 | 272.9 |
| microglia | SCING | 10.5 | 13.8 | 20.8 |
|  | ppcor | 4.4 | 3.9 | 4.2 |
|  | PIDC | 73.0 | 75.8 | 83.0 |
|  | GRNBOOST2 | 43.0 | 340.5 | 340.5 |
| oligodendrocytes | SCING | 10.3 | 13.5 | 20.7 |
|  | ppcor | 4.9 | 4.6 | 4.1 |
|  | PIDC | 74.5 | 77.3 | 86.0 |
|  | GRNBOOST2 | 13.8 | 29.0 | 71.3 |

**Table 2.** Average run time (seconds) of GRN building methods on variable number of cells for 1,000 genes across 10 iterations.

|  |  | 250 cells | 500 cells | 1,000 cells |
| --- | --- | --- | --- | --- |
| astrocytes | SCING | 9.9 | 11.0 | 10.6 |
|  | ppcor | 0.6 | 2.1 | 4.3 |
|  | PIDC | 0.2 | 19.6 | 74.1 |
|  | GRNBOOST2 | 9.8 | 14.8 | 33.3 |
| microglia | SCING | 10.1 | 11.0 | 10.5 |
|  | ppcor | 0.7 | 2.3 | 4.6 |
|  | PIDC | 9.2 | 18.7 | 73.1 |
|  | GRNBOOST2 | 10.2 | 15.3 | 45.8 |
| oligodendrocytes | SCING | 9.5 | 9.9 | 10.5 |
|  | ppcor | 0.52 | 2.2 | 4.7 |
|  | PIDC | 9.2 | 19.9 | 75.0 |
|  | GRNBOOST2 | 9.0 | 10.3 | 13.9 |

### SCING GRNs better predict downstream genes of perturbed genes in Perturb-seq

We tested whether GRNs from each approach can predict gene expression changes in downstream genes from gene knockdown treatments. Here, we used perturb-seq datasets, which enabled us to identify the effects of many perturbed genes in parallel. We utilize previously published datasets with THP-1, dendritic (DC), and K562 cells with 25, 24, and 21 genes perturbed, respectively^31,32^. The DC cells were split into lipopolysaccharide (LPS) stimulated and non-stimulated cells with perturbations targeting transcription factors (TFs) as two datasets, and the K562 cells were split into two datasets based on the genes initially perturbed (TFs or cell cycle related genes) in the Perturb-seq experiments. THP-1 cells contain perturbations targeting PD-L1 regulators.

We identified genes downstream of each perturbation through an elastic net regression framework to determine the effect of RNA guides on each gene while regressing out cell state^32^ (Methods). We compared GRNs generated from SCING to GRNs generated from GRNBOOST2, PIDC, and ppcor in predicting genes downstream of each target gene in each perturb-seq experiment. For any given network, we iterated through the downstream genes of an initially perturbed gene by the RNA guide and determined if the predicted downstream genes were significantly altered. We determined the true positive rate (TPR) and false positive rate (FPR) at each network depth to compute the area under the receiver operating characteristic (AUROC) curve, as well as the TPR at an FPR of 0.05. We examine TPR at FPR 0.05 to show perturbation prediction accuracy in a setting more relevant to biological analysis (controlling for FPR 0.05). We first examine the prediction performance of each GRN approach when building GRNs on datasets with cells removed that have zero expression of the target gene. Removing these cells mitigates performance effects from sparsity (SF2a). Since ppcor and PIDC produce undirected graphs, and GRNBOOST2 generally has bidirectional edges, we first evaluated SCING against the other methods without considering edge direction, showing a higher AUROC for SCING (Figure 2a). However, edge direction only affects prediction accuracy in TPR at FPR 0.05 for dc 3hr cells (Figure 2b). We additionally show SCING improves TPR at FPR of 0.05 (Figure 2c), and similarly, edge direction typically does not affect this metric (Figure 2d).

**Figure 2.**
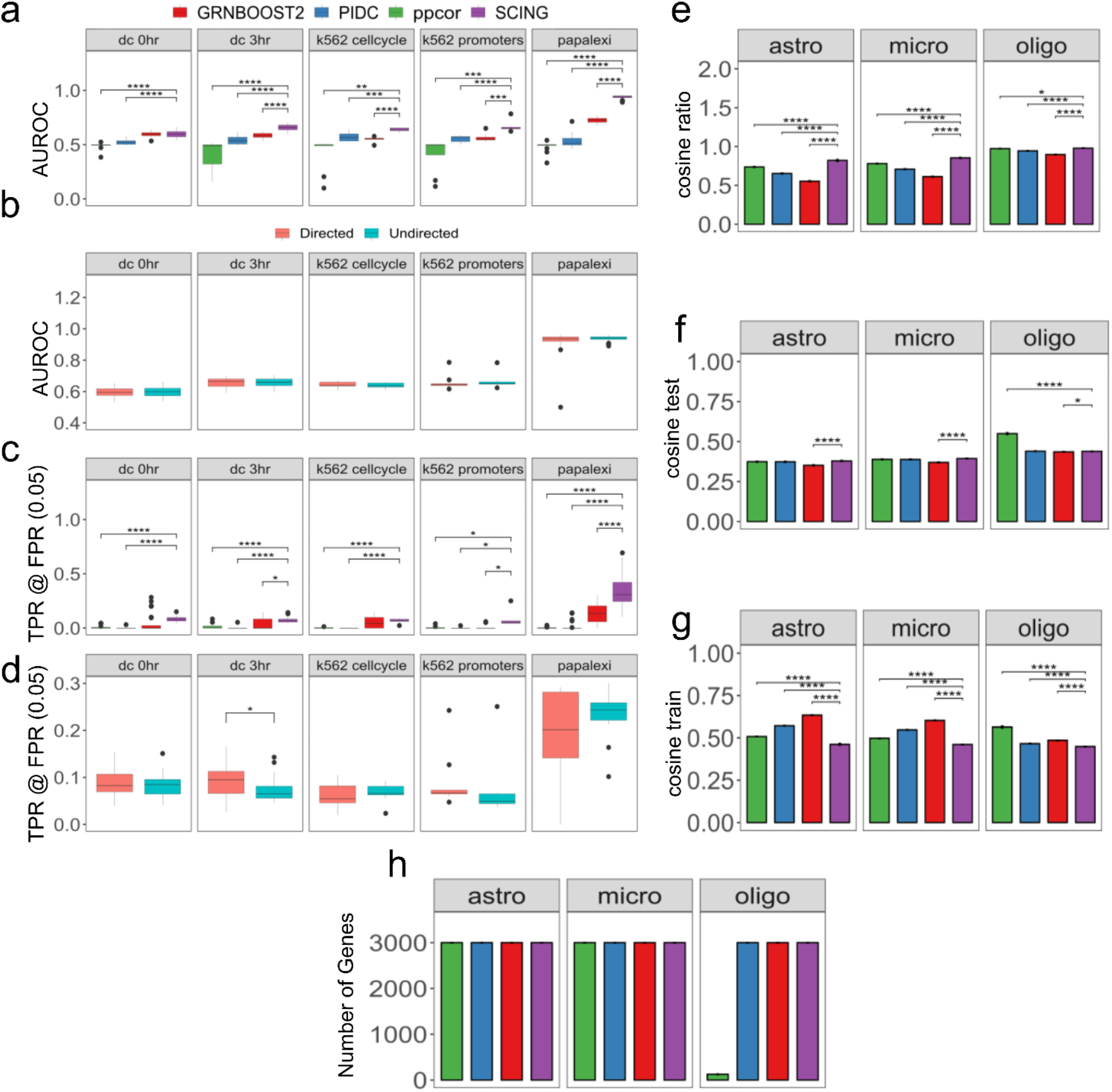
Performance evaluation. Predicted downstream affected genes of perturb-seq based perturbation in 5 datasets with GRNs built on cells with non-zero expression of the perturbation of interest. Area under receiver operator characteristic (AUROC) curve for prediction of downstream perturbations using undirected GRNs (a). AUROC for prediction of downstream perturbation on directed GRNs for SCING (b). True positive rate (TPR) at a false positive rate (FPR) of 0.05 for the prediction of downstream perturbations on undirected GRNs (c). TPR at FPR of 0.05 for the prediction of downstream perturbations on directed GRNs for SCING. (d) Measure of network overfitting by similarity of predicted gene expression and actual in held out data for astrocytes, microglia, and oligodendrocytes. Cosine similarity of predicted gene expression and actual in testing data show few differences between SCING and others (e). Cosine similarity of predicted gene expression and actual in training data shows other methods overfit to the training data (f). Ratio of test to train shows SCING does not overfit to the data as much as other approaches (g). (*: p < 0.05, **: p < 0.01, ***: p < 0.001, ****: p < 0.0001)

Across all perturb-seq datasets tested and when no cells were removed, SCING either outperformed, or met the performance of the other tools in both AUROC (SF2b) and TPR at FPR 0.05 (SF2d) and performance was minimally affected by edge direction (SF2ce). For AUROC, SCING outperforms ppcor across all datasets, PIDC across 4 datasets, and GRNBOOST2 across 2 datasets. For TPR at FPR 0.05, SCING outperforms ppcor and PIDC across 4 datasets and GRNBOOST2 across all datasets (SF2d). SCING performed best in the Papalexi et al. THP-1 dataset, although there is a large variance in both AUROC (SF2b) and TPR at FPR 0.05 (SF2d) for both ppcor and SCING.

Overall, SCING outperforms all other approaches at predicting perturbation effects in perturb-seq data when sparsity is adjusted (Figure 2) and outperforms select methods when all cells are used (SF2). Thus, we recommend removing cells with sparse gene expression when using SCING.

### SCING mitigates overfitting and builds more robust GRNs

Models often overfit to their data and fail to properly perform on new datasets. In the GRN context, we aimed to identify connections and networks that are able to capture biological variation rather than sample or batch specific effects. To test the performance of GRN inference approaches and their ability to capture robust biological signals, we tested the ability of a model trained on parents of each gene in training data to predict the gene expression of the downstream target gene in testing data. We split the scRNAseq data from control human prefrontal cortex^27^ into training and testing sets (Methods). First, we built GRNs on the training data from oligodendrocytes, astrocytes, and microglia using SCING, ppcor, PIDC, and GRNBOOST2. Subsequently, we trained gradient boosting regressors for each gene based on the parents in a given network using the training data. The trained regressors were then used to predict the gene expression of cells in testing data based on the expression of the parent genes in those cells. We evaluated the performance of each GRN approach by averaging the cosine similarity score over all downstream genes that have parents in the network. This process was repeated for 10 replicates on random subsamples of 3,000 genes to reduce runtime.

To measure overfitting, we used the cosine similarity score (ratio between the test and training sets) with a higher ratio indicating lower overfitting. We found that SCING GRNs had less overfitting than the other approaches (Figure 2e). In terms of performance in the test sets, SCING performed similarly to ppcor and PIDC, outperforming GRNBOOST2 (Figure 2f). On training data, GRNs from ppcor, PIDC, and GRNBOOST2 had higher cosine similarity scores compared to SCING, reflecting overfitting on the training data by the other methods (Figure 2g). We noted that the number of genes in the resulting network to be very low in the ppcor oligodendrocyte network (Figure 2h), which likely affected the results of ppcor as evaluated by the cosine similarity measure here. These results support the robustness of SCING GRNs, highlighting its ability to identify a robust GRN without overfitting or sacrificing performance.

### SCING fits scale free model, and shows edge consistency

As another measure of GRN quality and performance, we compared GRNs generated by each method by various standard network metrics (scale-free network fit, number of edges, number of genes, and betweenness centrality), as well as robustness of network edges between networks on 50/50 split datasets. We tested this on 10 replicates for each of the 3 cell types (oligodendrocyte, astrocyte, and microglia) in the scRNAseq data from control human prefrontal context, with 3,000 different genes randomly selected for each subsample of cells.

GRNs are thought to follow a scale-free network structure, in which there are few nodes with many connections, and many nodes with few connections^33^. We computed scale-free network structure through the R-squared coefficient of a linear regression model regressing on the log of each node’s degree and the log of the proportion of nodes with that given degree. We show that the R-squared value for SCING is significantly higher than that of the other methods, indicating SCING networks more closely follow a scale free network structure (Figure 3a). In a typical scale-free network plot of Iog10 node count vs Iog10 degree, we expect a power law distribution. However, we found that networks built on scRNAseq data have a parabolic distribution, with only the right half following a power law distribution whereas the left portion of the plot is driven by genes that were very sparse and likely do not fit typical distributions (Figure 3b). Therefore, we excluded the sparse genes from the regression calculation.

**Figure 3.**
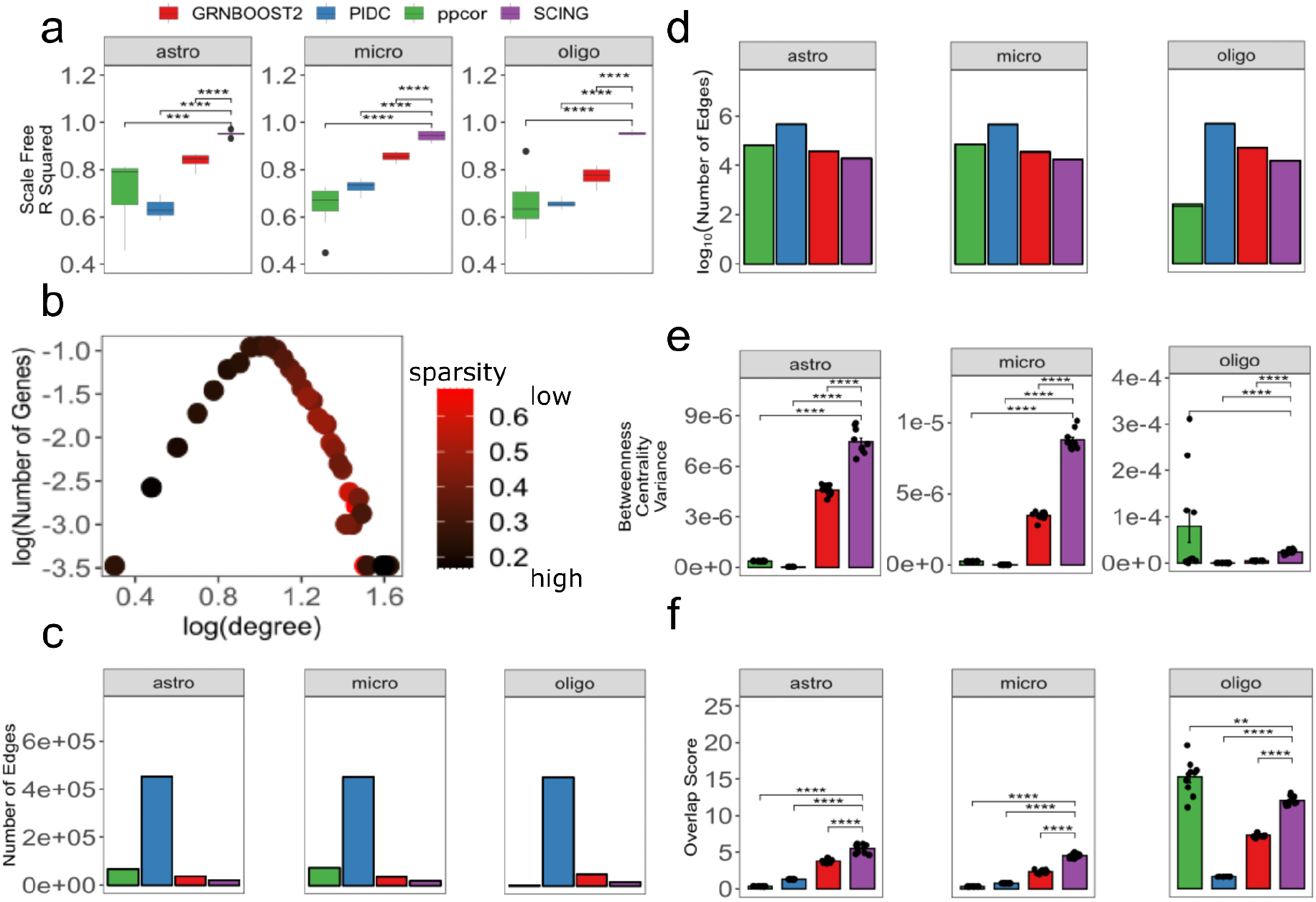
Network features and consistency. Descriptive features of networks across SCING and other approaches for 10 networks on astrocytes, microglia, and oligodendrocytes. Linear regression R-squared for log degree vs log count for goodness of fit metric of scale-free network (a). Example scatter plot of log degree vs log count with the average sparsity of genes in each dot. Brighter red indicates less sparse. This shows highly sparse genes tend to have lower degrees (b). Average number of edges (c) and log of the number of edges (d) for each method across all cell types. The variance of the betweenness centrality across nodes in each graph (e). The overlap score (number of overlapping edges/expected number of edges) in independent sets of cells (f). (*: p < 0.05, **: p < 0.01, ***: p < 0.001, ****: p < 0.0001)

Among all 4 methods tested, PIDC produced the largest networks (Figure 3cd), followed by ppcor, GRNBOOST2, and SCING. Since SCING aims to find robust edges, as expected, SCING networks have much fewer edges than the other approaches while keeping similar numbers of genes (Figure 2h). One exception is that the ppcor oligodendrocytes network contains only 214.8 edges on average compared to 14,545.8, 46,891.9, and 449,850 in SCING, GRNBOOST2, and PIDC, respectively. Many genes without regulatory edges were not included in the final ppcor network. The smaller ppcor oligodendrocyte network has implications for the betweenness centrality and edge overlap metrics, as follows.

Betweenness centrality is often used as another metric to determine the overall connectedness of a graph^34^. For a given node, betweenness centrality is the number of shortest paths that pass through that node, indicating how much information that node presents to the graph. High betweenness centrality indicates that a node conveys a lot of information to a given graph. We found that SCING networks generally have higher variance of betweenness for the nodes in the networks (Figure 3e). This indicates that some nodes are more centralized than others when compared to other approaches, again consistent with the scale-free network model.

To determine network consistency, we split each cell type into two groups of non-overlapping cells. We built networks for each dataset using all methods and calculated the fraction of total edges that overlap between the two networks. While larger networks tend to have more edge overlap, this also holds for larger random networks. We designed a normalized overlap score: the fraction of edges overlapped divided by the expected number of overlapping edges of a random network of the same size (Methods). When controlling for network size, SCING has significantly more overlap, or higher reproducibility, between networks of 50/50 split data than the other approaches (Figure 3f).

### SCING more accurately models disease subnetworks

To evaluate the performance of GRNs on disease modeling, we applied an approach developed by Huang et al^35^. Briefly, given a known disease gene set and a GRN, we evaluate the ability of the GRN to reach held out disease genes by starting from select disease genes in the network through random walks. We then compared the performance for each network to that of a random network in which the nodes follow similar degree characteristics to derive a performance gain measurement. Here, we selected known gene sets for 3 classes of diseases from DisGeNET (Immune, Metabolic, and Neuronal) (Supplementary Table 1) and obtained scRNAseq data for cell types from 3 tissues relevant to each disease class from the mouse cell atlas^26^ (bone marrow for immune diseases, brain for neuronal diseases, and liver for metabolic diseases) (Supplementary Table 2). First, to reduce the number of genes, we filtered the scRNAseq data by removing genes expressed in fewer than 5% of cells and added expressed disease genes from all DisGeNET disease gene sets. We built GRNs using each method and evaluated the performance gain over random networks on the disease gene sets. We found that across all tissues and all disease types, SCING outperformed all other approaches (Figure 4a).

**Figure 4.**
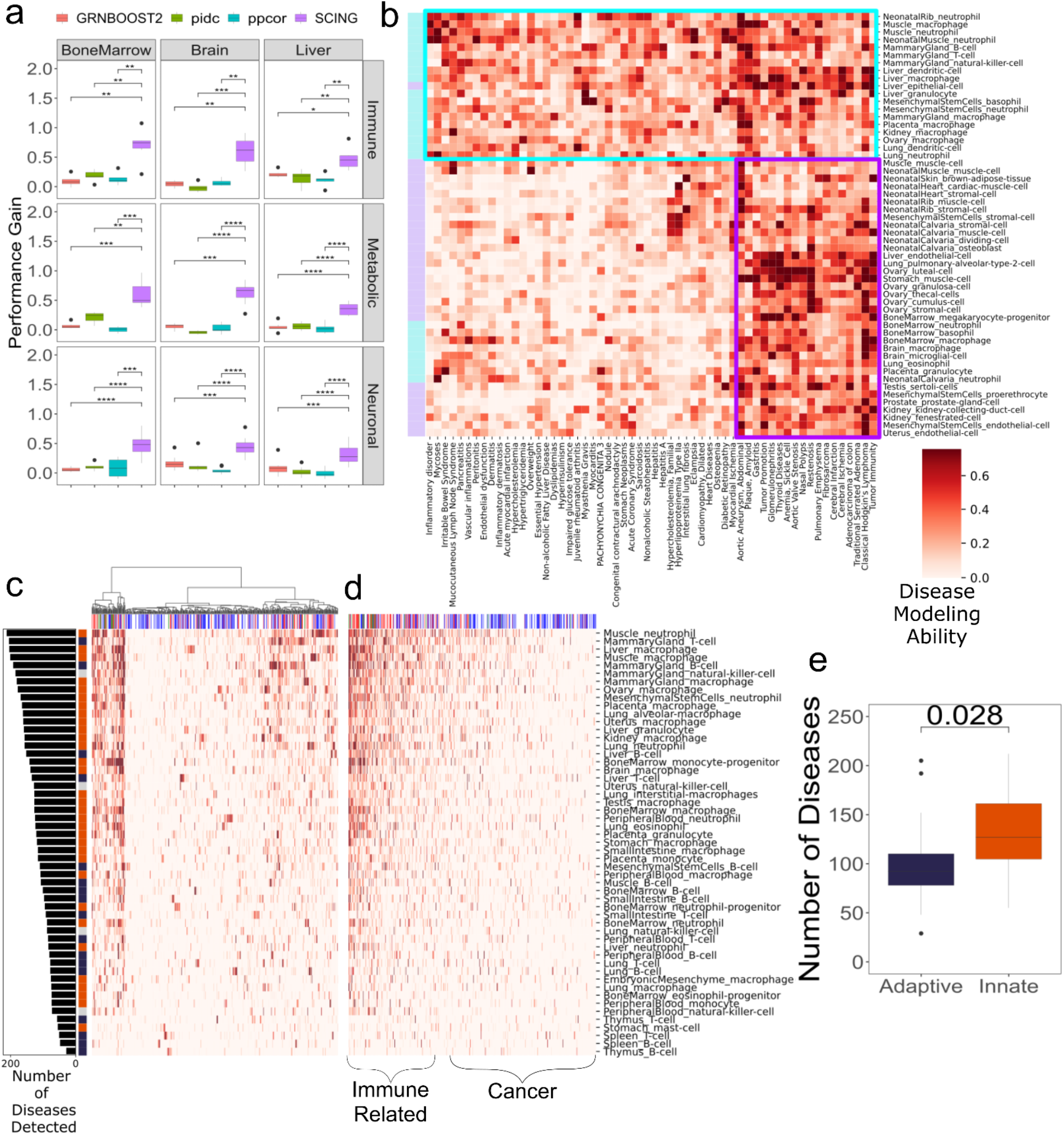
Application case 1: constructing and using SCING GRNs based on Mouse Cell Atlas scRNAseq datasets to interpret diseases. Performance of modeling disease subnetworks for DisGeNET gene sets related to the immune, metabolic, and neuronal diseases with GRNs built on bone marrow, brain, and liver cells, reveals SCING models disease subnetworks more accurately than other methods (a). Clustermap depicting GRNs built with SCING from immune cell types (light blue), model disease subnetworks from many different disease gene sets, while vascular cell types (purple) are more specific to vascular diseases (b). Cell types (rows) from the adaptive (blue) and innate (orange) immune systems, show variability in the number of diseases (columns) they model (>0.1). Clustermap shows diseases clustered with hierarchical clustering (c) and sorted by the number of cell types that can accurately model that disease subnetwork (d). Diseases are colored by disease category (immune related: red; cardiothoracic: green; cancer: blue; immune related cancer: purple), and cell types are colored by innate (orange), and adaptive immune system (dark blue). Innate immune system cell types better model disease subnetworks from more diseases (e). (*: p < 0.05, **: p < 0.01, ***: p < 0.001, ****: p < 0.0001)

### Application case 1: constructing SCING GRNs using Mouse Cell Atlas (MCA) scRNAseq datasets to interpret diseases

After establishing the performance of SCING GRNs using the various approaches described above, we established a SCING GRN resource for diverse cell types and tested the broader utility of SCING to produce biologically meaningful GRNs. To this end, we applied SCING to generate GRNs for all cell types with at least 100 cells in all tissues of the MCA. We constructed a total of 273 cell-type specific networks, across 33 tissues and 106 cell types. To identify which GRN informs on which disease, we applied the above random walk approach from Huang et al. and summarized the results in (SF3abc, Supplementary Table 3). We found clusters of cell type GRNs defined by DisGeNET diseases that had similar patterns (SF3a, Figure 4b). Some disease genes can be modeled well using GRNs from numerous cell types (SF3a) whereas others are more cell type or tissue specific (SF3b). Additionally, some cell type GRNs are able to model a broad range of diseases (SF3ac). We found that immune cell type (light blue squares in Figure 4b) GRNs can model a wide range of diseases, whereas non-immune cell type GRNs (light purple squares in Figure 4b) are more specific to vasculature related diseases.

We further explored the dynamics of GRNs of immune cell types across all diseases in DisGeNET. We clustered the cell types in the performance gain matrix with only immune cell types included, and sorted the GRNs by the number of diseases they can accurately model (Figure 4cd, SF4ab). We noticed that cell types of the innate immune system can model a broader range of diseases than those of the adaptive immune system^36^ (Figure 4e).

Our SCING cell type GRNs resource and the above patterns of relationships between cell type GRNs and diseases support the utility of the SCING cell type GRN in disease interpretation. The networks can be accessed at https://github.com/XiaYangLabOrg/SCING to facilitate further biological mining of complex diseases.

### Application case 2: Using SCING GRNs to interpret Alzheimer’s disease (AD)

We next applied SCING to a single nuclei RNAseq (snRNAseq) dataset from Morabito et al. that examined human prefrontal cortex samples from AD and control patients to evaluate the applicability of SCING GRNs in understanding AD pathogenesis^27^. We focused on microglia, due to their strong implication in AD^37,38^ and a better current understanding of the genes and biological pathways in microglia in AD, to demonstrate that SCING GRNs can retrieve known biology.

The SCING microglia GRN contained 10,159 genes and 63,056 edges. Using the Leiden clustering algorithm^39^, we partitioned the SCING microglia GRN into 21 network modules. Next, we summarized module-level expression for each cell using the AUCell method from SCENIC on the partitioned GRN modules^19^. When cells were clustered based on the raw gene expression values, as is typical with human samples, cells from individual samples clustered together (Figure 5a), making it difficult to isolate sample heterogeneity from biological variability. However, when using SCING module expression to cluster cells, the sample, batch, and RNA quality effects were mitigated (Figure 5ab). In contrast, biologically relevant variation, such as sex, AD diagnosis, and mitochondrial fraction were better retained (Figure 5cd). In the UMAP control cells tend to localize to the right side, while cells from females tend to localize to the top part. These results suggest that SCING GRNs have intrinsic ability to correct for non-biological variations.

**Figure 5.**
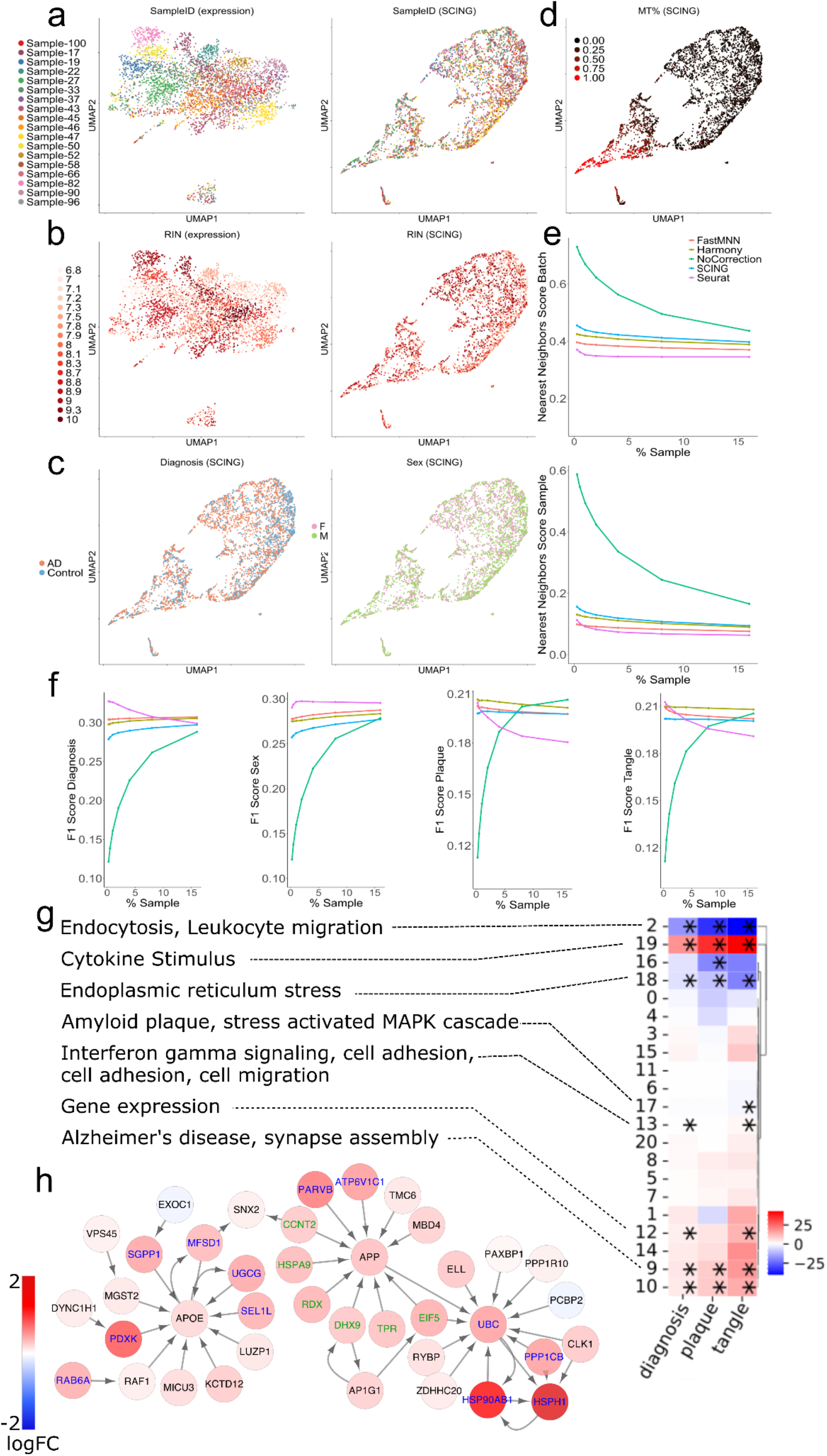
Application case 2: Using SCING GRNs to interpret Alzheimer’s disease (AD). UMAP representation of scRNAseq data shows sample specific differences when operating on gene expression space (left panel). Dimensionality reduction on SCING module embeddings removes sample specific effects (right panel). SCING removes RNA quality effects on gene expression clustering (b). Clustering on SCING modules, keeps biologically relevant features such as AD status (left panel) and sex (right panel) (c), as well as mitochondrial fraction (d). SCING corrects batch (top) and sample (bottom) specific effects based on the fraction of neighbors with the same batch or sample status (lower is better) for a given cell (e). This was performed at different fractions of the cells used as neighbors for a given cell. Dedicated batch correction techniques have better batch correction. F1 score of nearest neighbor score with batch and sample effects corrected (higher is better), shows SCING corrects sample, and batch effects while retaining biologically relevant features (f). Heatmap showing coefficients of linear regression of diagnosis, plaque, and tangle status, on module expression, while regressing out sex (g). Pathway annotations for significant (*: FDR < 0.05) modules are provided. Subnetwork reveals collocalization of canonical Alzheimer’s genes and subnetworks such as APOE and APP (h). Colors indicate the linear regression coefficients between a phenotype and module expression.

To quantitatively evaluate how well SCING GRNs can be used for batch effect correction and biological preservation, we compared SCING GRNs with commonly used batch correction methods, such as FastMNN^40^, Harmony^41^, and Seurat^42^, chosen based on their better performance in previous benchmarking studies^43^. We first performed dimension reduction and clustering of cells based on corrected data from each batch correction method. For SCING, the values used were SCING GRN module AUCell scores. We took each cell, determined how many neighbors in the PC space had the same annotation of interest (sample, batch, diagnosis, etc.), and then scored each batch correction approach by the fraction of cells that had the same annotation. We removed batch and sample specific effects using the F1 score (Methods)^43^. We found that the SCING GRN module based dimensionality reduction carried the ability to correct for batch effect and retain biological information (Figure 5ef) in a similar manner to dedicated batch correction methods such as FastMNN^40^, Harmony^41^, and Seurat^42^. Although SCING GRN based batch correction was not as optimized as the dedicated batch correction methods, SCING is unique in that each GRN module has direct biological interpretability, since each SCING module can be associated with phenotypic traits and annotated with pathways, as described below.

We identified SCING GRN modules associated with AD diagnosis, plaque stage, and tangle stage through linear regression of the phenotypic traits and each module’s expression across cells, while regressing out sex specific differences. We found that ~43% of modules were significantly (FDR < 0.05) associated with at least one trait and ~24% of modules were significantly associated with all three traits. To examine the biological interpretation of these modules, we performed pathway enrichment on the genes in each module utilizing the GO biological process, DisGeNET, Reactome, BioCarta, and KEGG knowledge bases. We found that 78% of the significantly trait-associated modules were significantly enriched for biological pathways (Figure 5g, Supplementary Table 4). These modules recapitulated pathways related to Alzheimer’s disease (module 9), immune processes (modules 0, 2, 9, 13, and 19), cytokine triggered gene expression (modules 12, 18, 19), and endocytosis (modules 2, 9, and 13). These are expected perturbed pathways for microglia in AD^37^.

We dug into the vesicle mediated transport pathway from module 2 and visualized the network with the differential gene expression of each gene (SF5). This microglia pathway is important in AD^44^. Additionally, we found the APOE and APP subnetwork within the vesicle mediated transport pathway which are among the most significant genetic risk factors for AD (Figure 5h)^45–47^.

Therefore, our results demonstrate that SCING GRNs can correct for batch effects intrinsic to scRNAseq studies and can recapitulate known cell type specific genes, pathways, and network connections.

### Application case 3: Using SCING to model GRNs based on 10x Genomics Visium spatial transcriptomics data to interpret AD

To evaluate the applicability of SCING beyond scRNAseq and snRNAseq, we next applied SCING to spatial transcriptomics data from Chen et al.^28^ as a new approach for spatial transcriptomics analysis. We built a SCING GRN on all spots in the visium data to obtain a global GRN with 128,720 edges across 15,432 genes. Additionally, Chen et al. profiled beta amyloid plaque with immunohistochemistry, and we included this protein expression value in the SCING GRN, highlighting the possibility of constructing multi-omics networks using SCING.

We partitioned the genes in the resulting SCING GRN with the leiden graph partition algorithm into 33 modules and performed the AUCell score from SCENIC^19^ to obtain module specific expression for each spot and annotated the enriched pathways for each module (Supplementary Table 5). We found module specific expression in the mouse brain subregions (Figure 6ab, SF6), based on clustering of the average module expression across spots in each brain subregion (SF7). We found cortical subregions to cluster together, as well as the thalamus and hypothalamus (SF7), based on GRN module expression patterns. We also identified modules more specifically expressed in the cortex and hippocampus (CS, HP) (module 12), or the fiber tract, thalamus, and hypothalamus (BS) (module 14) (Figure 6ab). Module 12 was highly enriched for genes involved in neuronal system, axonogenesis, and chemical synapse, which might reflect the dynamic status of hippocampus and cortex neurons for memory formation and cognitive function. In contrast, module 14 was enriched in genes involved in myelination^48^ and blood-brain barrier^49^, consistent with the high enrichment of oligodendrocyte populations in the fiber tracts, and indicated blood-brain barrier changes in the thalamus. We also found modules that separate more similar subregions from one another such as module 27 (enriched for axonogenesis) and 21 (enriched for calcium ion transport) expressed much higher in the thalamus than in the hypothalamus (SF7). By contrast, modules, such as module 5 (enriched for chromatin organization and peroxisomal lipid metabolism) and module 19 (enriched for ribosomal biogenesis and protein processing), are much less specific to subregions (SF6, SF7). These are general cellular functions that are expected to have broad expression across the brain.

**Figure 6.**
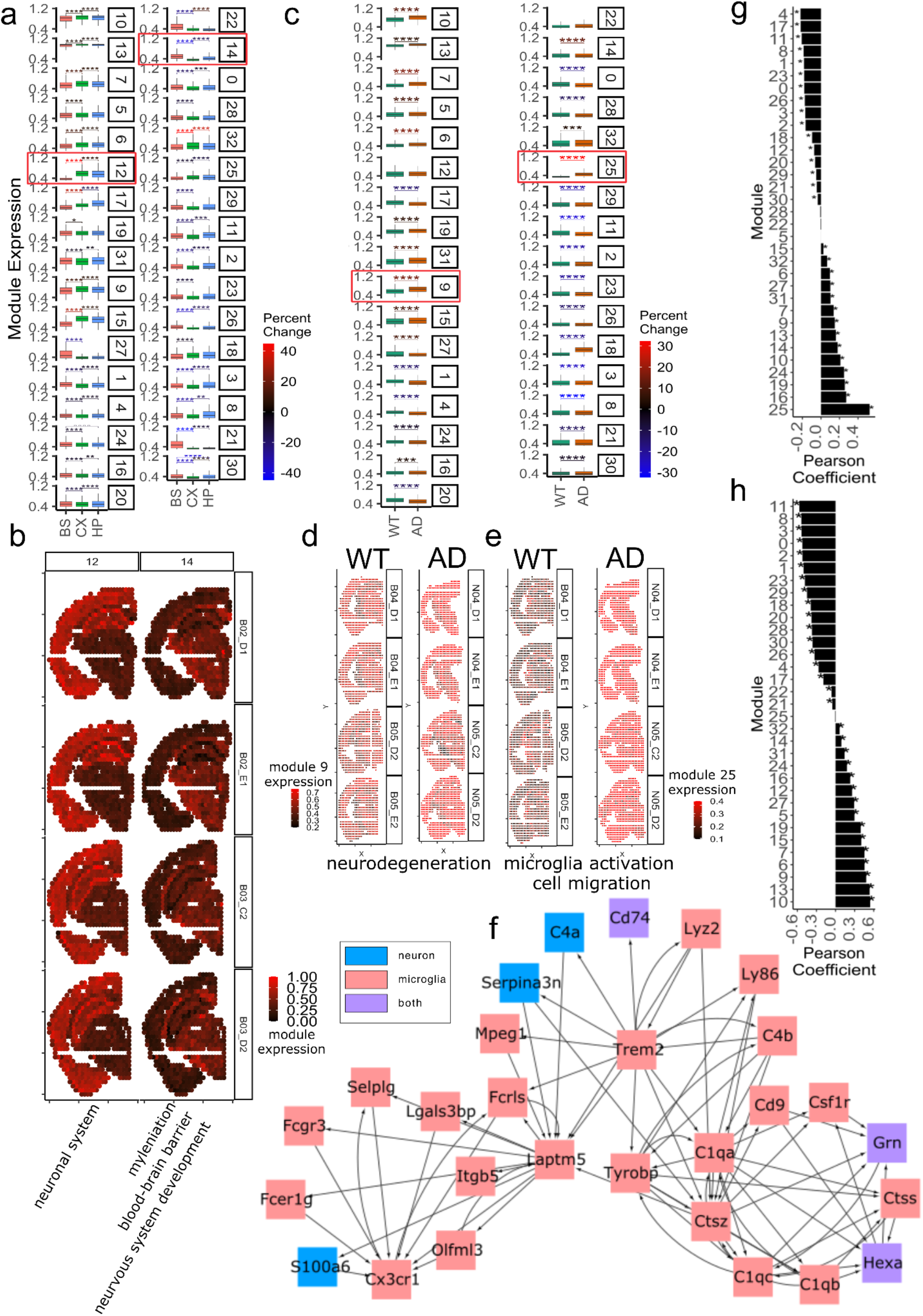
Application case 3: Using SCING to model GRNs based on 10x Genomics Visium spatial transcriptomics data to interpret AD. Boxplots show regional specificity of specific modules, namely module 12 and 14 (red boxes) (a). Scale bar shows the t-test statistic of the test between regions. Regional specificity of module 12 (hippocampus and cortex) and module 14 (hypothalamus, thalamus, and fiber tract), visualized on brain samples of 3 month old WT mice (b). Module association in AD vs WT mice (c). Scale bar shows the t-test statistic of the test between mouse groups. Visualization of module 9 in 18 month old WT and AD mice (d). Visualization of module 25 in 18 month old WT and AD mice (e). Module 25 subnetwork of Trem2 and complement proteins shows microglial association of module 25 (f). Full subnetwork in supplement 9. Nodes are colored by marker gene status of neuronal (blue), microglial (red), or both (purple), as determined from the Allen Brain Atlas. Cross cell type communication edges seen between red and blue. Pearson correlation between module expression and plaque (g) and age (h), shows with significance (*: p<0.05) Subpanel a,c: (*: p < 0.05, **: p < 0.01, ***: p < 0.001, ****: p < 0.0001)

We found many modules to be AD associated (Figure 6c), in particular, module 9 (enriched for neurodegeneration) (Figure 6d) and module 25 (enriched for microglial activation, lysosome, and cell migration) (Figure 6e). We further explored these subnetworks and found most of the genes in module 9 to be neuronal marker genes, while most of the genes in module 25 to be microglial marker genes (SF 9). The module 25 subnetwork to contain the *Trem2* and *C1q* subnetworks, highly profiled in AD microglial cells (Figure 6f). Interestingly, we found a cross-module edge and several cross-cell-type edges, likely revealing intercellular communications.

We also identified all but two modules that were significantly correlated with amyloid beta plaque (Figure 6g). Modules most highly correlated with plaque were module 25 (enriched for immune function), module 19 (enriched for nonsense mediated decay, and infectious disease), and module 9 (enriched for neurodegeneration), which were all also associated with AD (Figure 6c).

We found that amyloid plaque staining could be partitioned based on the expression of module 10 in the SCING GRN (SF8), which had significantly higher expression in AD mice than in WT (Figure 6c), and is also quantitatively correlated with plaque (Figure 6g). Module 10 contains genes highly related to metabolism, neurodegenerative diseases, and immune function (Supplementary Table 5). We show the subnetwork for the amyloid beta plaque in module 10 (SF9). Additionally, module 25 (enriched for microglial activation, lysosome, and cell migration) was also highly associated with plaque (Figure 6g), as expected in AD pathogenesis.

In addition to plaque association, we explored the age related modules through pearson correlation of age (6, 12, and 18 months) and module expression in WT samples. We found most modules to be age related (Figure 6h); however, some modules such as module 21 (enriched for muscle contraction and calcium ion transport) have a lower correlation coefficient. The top 2 positively correlated modules with age, module 10 (subnetwork with amyloid protein) and 13, have age related pathways. Module 10 is involved in protein degradation, neurodegeneration, and cell cycle, and module 13 is involved in metabolism, cellular response to stress, and immune system (Figure 6h, Supplementary Table 5).

We found certain modules such as module 30 (enriched for neuropeptide signaling) to be spatially variable (SF7) and associated with AD (Figure 6b). Based on the locations from the Allen Brain Atlas^50^ (SF10a), we find module 30 to be more specific to the hypothalamus than other regions of the BS (SF10b). We also find that within the hypothalamus, module 30 expression is higher in 18 month old AD mice compared to the WT mice (SF10c). Hypothalamic alterations have been observed in the hypothalamus in AD development^51^. The module 30 subnetwork (SF10d) shows key hypothalamic neuropeptides, such as Pome^52^ and Pnoc^53^ which is consistent with their enrichment in the hypothalamus.

Our applications of SCING to spatial transcriptomics data demonstrate its broader utility beyond scRNAseq/snRNAseq and revealed spatial network patterns of AD.

## Discussion

Single cell multiomics has become a powerful tool for identifying regulatory interactions between genes, but the performance of existing tools is limited in both accuracy and scalability^13^. Here, we present SCING, a gradient boosting, bagging, and conditional mutual information based approach for efficiently extracting robust GRNs on full transcriptomes for individual cell types. We validate our networks using a novel perturb-seq based approach, held-out data prediction, and established network characteristic metrics (network size, network overlap, scale-free network, and betweenness centrality) to determine performance and network features against other existing tools. SCING not only offers robust and accurate GRN inference and improved gene coverage and speed compared to previous approaches^9,18^, but also versatile GRN inference with scRNAseq, snRNAseq, and spatial transcriptomics data. Using various application examples, we show that SCING infers robust GRNs that inform on cell type specific genes and pathways underlying pathophysiology while simultaneously removing non-biological signals from data quality, sample, and batch effects through gene regulatory module detection and functional annotation. We also provide a comprehensive SCING GRN resource for 106 cell types across 33 tissues using data from MCA to facilitate future applications of single cell GRNs in our understanding of pathophysiology.

SCING efficiently identifies robust networks using supercells, a bagging approach, and mutual information based edge pruning, to remove redundant edges in the network. The supercell and reduction in potential edges make the bagging approach possible by removing computational time for each GRN. GRNBOOST2 uses a similar framework as SCING (gradient boosting regression) but overfits the data by generating too many edges to be interpretable, which are also undirected, making them less useful for biological interpretation. Meanwhile, ppcor and PIDC use partial correlation and partial information decomposition approaches, which are more accurately described as measures of coexpression, rather than gene regulation. In contrast, the directed graphs built with SCING show better perturbation prediction and consistency across replicates.

Validation of GRN inference tools has remained challenging^13^. Our novel perturb-seq based approach provides a unique way to determine GRN accuracy. Prediction of perturbed genes is a very powerful aspect of GRN construction, and SCING stands out above all other methods in this regard^54^.

We demonstrate that SCING networks are applicable to scRNAseq, snRNAseq, and spatial transcriptomics data. Our network-based module expression provides batch corrected, biologically annotated expression values for each cell that can be directly used for disease modeling (Figure 4), phenotypic correlation (Figure 5), and spatially resolved analysis (Figure 6) to boost our ability to interpret single cell data.

At the intersection of single cell omics and complex diseases, SCING provides sparse but robust, directional, and interpretable GRN models for understanding biological systems and how they change through pathogenesis. GRNs can be analyzed to identify and predict perturbed subnetworks, and as a result, be used to investigate key drivers of disease^55^. Identifying key drivers of disease by teasing apart biology and technical variation from high throughput, high dimensional datasets will lead to more successful drug and perturbation target identification, as well as robust drug development^56–58^.

We note that SCING is currently tested to infer GRNs based on individual scRNAseq, snRNAseq, and spatial transcriptomics data. Other types of omic information such as scATACseq, scHi-C, or cell type specific trans-eQTL information can be included in SCING to further inform on regulatory structure to refine and improve on GRNs^9,59–62^. Information from multiple data types will become an integral part of the systems biology, and future efforts to properly model multiomics data simultaneously to inform on complex disease are warranted.

## Methods

### SCING method overview

To reduce the challenges from data sparsity from single cell omics, as well as reduce computational time, we first used supercells which combine gene expression data from subsets of cells sharing similar transcriptome patterns. To improve the robustness of GRNs, we built GRNs on subsamples before merging the networks, keeping edges that appear in at least 20% of networks. Lastly, we removed cycles and bidirectional edges where one direction was >25% stronger than the other direction and pruned the network using conditional mutual information to reduce redundancy. These steps are described in more detail below. To benchmark the performance of SCING, we selected three existing methods, namely GRNBOOST2, ppcor, and PIDCm with default parameters. These methods were selected based on previous benchmarking studies where superior performance of these methods were supported^13^.

### Supercell construction

For each dataset, we first normalized the data for a total count number of 10,000 per cell^63^. We then took the log of the gene expression values, identifying and subsetting to the top 2000 highly variable genes. Data were then centered and scaled and used to compute the nearest neighbor embedding with 10 neighbors. We then used the Leiden graph partitioning algorithm to separate cells into groups. The leiden resolution is determined by the user input specifying the final number of supercells. Here, we used 500 supercells which balances runtime, with dataset summarization. We then merged each group of cells into a supercell by averaging the gene expression within each group.

### GRN inference in SCING

We normalized the total number of counts in each cell to 10,000 and took the natural log of the gene expression. We removed genes not expressed in any cells and any duplicate genes. We transposed, centered, and scaled the data before running principal component analysis (PCA). This provides us with low dimensional embeddings for each gene. A nearest neighbor algorithm from scikit-learn^64^ was used to find the nearest neighbors of each gene. The potential regulatory relationship between genes was limited to the 100 nearest neighbors. For each gene, we trained a gradient boosting regressor^18^ to predict the expression based on its nearest neighbors. For each gradient boosting regressor, we used 500 estimators, max depth of 3, learning rate of 0.01, 90 percent subsample, the square root of the total number of features as max features, and an early stop window of 25 trees, if the regressor was no longer showing improved performance^18^.

### SCING parameter selection

Parameter selection was used to retain biological accuracy while limiting computational cost. We determined that using 500 supercells and 100 GRNs per merged network balanced computational resourcefulness with robustness. If computational cost is not an issue, supercells are not necessary and more GRNs can be built as intermediates. Since GRNs are typically built within a single cell type, we use 10 principal components (PCs) for determining gene covariance. Typically in scRNAseq analysis more PCs are used, but there is less variation overall within one cell type. We determined 100 neighbors to be chosen for each gene, again balancing computational cost.

### Merging GRNs from data subsamples

For each network from subsampled data, we kept the top 10 percent of edges based on the edge weight, as well as the top 3 edges for each downstream gene and edges that appeared in greater than 20% of all networks. We also removed reversed edges if the edge with a higher weight was at least 25% stronger than that of the reverse direction. Otherwise, we kept the edge bidirectional. We removed cycles in the graph by removing the edge with the lowest edge weight. We removed triads in the network based on the significance of the conditional mutual information. The p-value of the conditional mutual information is based on the chi-squared distribution^65^. If an edge between two genes is not statistically significant given a parent of both of the genes, then the edge is removed.

### Other GRN methods

We benchmarked SCING against GRNBOOST2, ppcor, and PIDC. Default parameters were used for all existing approaches unless otherwise specified.

For GRNBOOST2^18^, we ran this approach by predicting the expression of all genes from all other genes. We then took the top 10% of edges to reduce the number of edges with extremely low importance (e.g. 10^-17^).

For ppcor^16^, due to the sparse non-linear nature of scRNAseq connections, we ran the approach using spearman correlation. We only kept edges with a Benjamini-Hochberg FDR < 0.05.

For PIDC^17^, according to their tutorial, we used a threshold of 0.1, to keep the top 10% of highest scoring edges.

### Datasets

For the perturb-seq validation, we used datasets from Dixit et al.^32^ and Papalexi et al.^31^. Dixit et al. has 24 transcription factors perturbed in dendritic cells and 25 cell cycle genes targeted in K562 cells, while Papalexi et al. has 25 PD-L1 regulators perturbed in THP-1 cells.

For the train-test split and network consistency assessment, we used the human AD snRNAseq data from Morabito et al.^27^. This adds another slightly different data type from the scRNAseq in the perturb-seq and MCAand is later used for biological application with microglial cells in AD.

We used the mouse cell atlas^26^ scRNAseq database, since it has a large number of cell types (106 cell types) across numerous tissues (33 tissues), to test 446 disease associations with the random walk approach from Huang et al.^35^ This additionally provides a resource of GRNs throughout cell types of the entire mouse.

Finally, to test the applicability of SCING on spatial transcriptomics data, we used the mouse AD dataset from Chen et al.^28^ This dataset contains AD and WT mice from various age groups (3, 6, 12, 18 months), in addition to amyloid beta plaque staining.

### Computation of time requirements

We determined the run time to build a GRN from each approach on subsets of cells and genes with varying cell numbers and gene numbers. All tests were performed on a ryzen 9 3900X 12-Core processor with 64Gb RAM. We determined the speed on 10 iterations of randomly selected genes (1000, 2000, 4000) with 1000 cells each, and on randomly selected cells (250, 500, 1000) with 1000 genes each.

### Overview of network robustness evaluation based on Perturb-seq datasets

Briefly, to determine the accuracy of the GRNs from each approach with perturb-seq data, we first identified significantly altered genes downstream of each guide RNA perturbation through an elastic net regression approach^32^, as detailed below. We then determine the accuracy of a given network by identifying the true positive rate (TPR) and false positive rate (FPR) at each depth in the network. The AUROC and TPR at FPR 0.05 were determined for each network on a given perturbation. More details on each step are below.

### Computation of guide RNA perturbation coefficients

First, we downloaded the perturb-seq data from Papalexi et al.^31^ and Dixit et al^32^. We then followed the steps as described by Dixit et al^32^ to compute guide RNA perturbation coefficients, which indicate the effects of a specific guide RNA (single perturbation) on other genes. As cell state can affect gene perturbation efficiency, to determine cell states and remove state specific perturbations, we clustered the non-perturbed cells in each dataset through Leiden clustering. The leiden resolution was determined by identifying unique subclusters in the data (Table 3). We subset the data to highly variable genes (min_mean=0.0125, max_mean=3, min_disp=0.5). We centered and scaled the data and performed PCA to get the first 50 PCs. We then trained a linear support vector machine (C=1) on the 50 PCs of the data and determined the probability of each cell in the dataset being from each state. We used these continuous state probabilities in our regression equation to regress out state specific effects on gene expression. To identify the perturbation effect of a given guide RNA on genes other than the target gene, we utilized elastic net regression (I1_ratio=0.5, alpha=0.0005)^32^. We fit the elastic net model to predict the gene expression of all genes from the binary matrix determined by the guide sequenced in each cell, combined with the continuous state values determined for each cell. To remove the effect of synergistic perturbations, we removed cells with multiple perturbations. We determined each guide’s perturbation effect on a given gene by the regression coefficient.

**Table 3.** Hyperparameters for perturb-seq clustering

| Dataset | Resolution |
| --- | --- |
| DC 0hr | 1.2 |
| DC 3hr | 0.7 |
| K562 cell cycle | 1.0 |
| K562 promoters | 1.0 |
| Papalexi | 1.2 |

### Determination of significant perturbation effects

To determine the significance of a given perturbation coefficient, we employed a permutation test as in Dixit et al^32^. For each guide, we permuted the vector of perturbations to randomize which cells received the given perturbation of interest. The elastic net regression model was trained with the same hyperparameters to determine the coefficients of perturbation. This approach was repeated 100 times to generate a null distribution of the perturbation effect of a given guide on each gene. The p-value was calculated as the fraction of null coefficients that were greater than or less than the true coefficient, determined by the sign of the coefficient. Significant perturbations were determined at a false discovery rate of 0.05 using the Benjamini-Hochberg procedure. This permutation approach was repeated for each guide RNA. A gene was determined as a downstream perturbation if at least one guide had a significant perturbation for the given gene.

### Selection of genes and cells for perturb-seq networks

To reduce computational cost and enable network building for all approaches, we first took the top 3,000 highly variable genes using the variance stabilizing transform method^66^, including the differentially perturbed genes. We built two networks for each dataset, one using all cells and the other using only cells with non-zero expression of the gene of interest.

### Evaluation of perturbation predictions

We built networks for each perturbed gene separately using SCING, GRNBOOST2, ppcor, and PIDC. Starting from the perturbed gene of interest, at each depth in the network, we determined the TPR and FPR based on the perturbed genes computed above. This gives a TPR vs FPR graph, from which an AUROC was computed. For each perturbed gene, we calculated the AUROC and the TPR at a FPR of 0.05.

### Network robustness evaluation based on training and held out testing data

We performed a train test split (75/25) on the dataset from Morabito et al on each random subsample of 3000 genes. We built GRNs on the training data and trained a gradient boosting regressor for each gene based on the predicted regulatory parents. We used the trained gradient boosting regressor to predict the expression of all genes in the test dataset and evaluated the performance based on the cosine similarity metric. We performed this on the training and testing data separately and computed the test to train ratio of the cosine similarity, with a smaller test to train ratio indicating potential overfitting of the training data.

### Computation of network characteristics

We built GRNs on oligodendrocytes, astrocytes, and microglia from snRNAseq data from Morabito et al^27^. For each cell type, we randomly selected 3000 genes (reduce computational time of methods) for each sample and generated 10 GRNs, in which 3000 genes were randomly subsampled from the full transcriptome. To compute scale-free network characteristics for each network, we fit a linear regression model on the log of each node degree with the log of the proportion of nodes at each degree. We removed low degree data points that are an artifact of scRNAseq sparsity. We also characterized each network by the number of edges in the network, number of genes remaining in the resulting network, and the mean betweenness centrality of nodes across the network.

### Computation of network overlap

We used the Morabito et al datasets described above and split each dataset in half and generated GRNs on each subset of cells. We checked the overlapping edges between the two networks and normalized for the expected number of overlapping edges based on the number of total edges in each network and the hypergeometric distribution. The overlap score measured the fraction of overlap between the two networks, divided by the expected number of overlapping edges.

### Assessment of disease subnetwork retrieval of GRNs

We utilized a random walk approach from Huang et al^35^ to determine the ability for GRNs from different methods to accurately model disease gene subnetworks. This approach provides a biologically relevant benchmarking approach to determine a GRNs ability to model disease subnetworks. Briefly, the approach splits a known disease gene set into two groups, to attempt to reach the held out gene set starting from the selected disease genes through random walks. An improvement score is computed by calculating the z-score for a given network relative to 50 degree-preserved randomized networks.

We built networks from the MCA on immune cells from bone marrow, neurons from the brain, and hepatocytes from the liver. To accommodate less efficient tools, we subsetted the transcriptome to genes that are expressed in more than 5% of the cells in the dataset. We used the method from Huang et al. using relevant immune, neuronal, and metabolic disease gene sets from DisGeNET. We kept these genes with >5% percent expression and included the genes from the disease gene sets. We determined performance of each subnetwork based on the improved performance compared to the random network distribution.

### Application of SCING to construct GRNs for all MCA cell types and assessment of network relevance to all DisGeNET disease gene sets

We applied SCING to all cell types for all tissues in the MCA and utilized the approach from Huang et al. to determine the ability of each network to accurately model each disease gene set. We clustered the disease gene sets and cell types using hierarchical clustering with complete linkage. We determined the number of disease sets accurately modeled by each cell type based on a performance gain of at least 0.1. We subsequently computed the number of cell types that can accurately model each disease set. To compare the number of diseases modeled by cell types from the adaptive and innate immune system on tissue relevant subsets of the DisGeNET diseases, we performed a t-test between the distributions of the number of disease gene sets each cell type can accurately model.

### Biological application to microglia in Alzheimer’s disease patients

We built a SCING network for the microglia on the genes expressed in at least 2.5 percent of cells in the Morabito et al. dataset^27^. For the SCING pipeline, we used 500 supercells, 70 percent of cells in each subsample, 100 neighbors, 10 PCs, and 100 subsamples. We utilized the Leiden graph partitioning algorithm to divide genes in the resulting GRNs into modules. We performed Leiden clustering at different resolutions and performed pathway enrichment analysis on the modules using the enrichr^67^ R package, using the GO biological process, DisGeNET, Reactome, BioCarta, and KEGG knowledge bases. We selected the resolution (0.0011) that had the highest fraction of modules annotated for between 20 and 50 modules per network. This avoids clustering too many modules with few genes while maintaining enough separate modules to have biological interpretation. We used the AUCell method from the SCENIC workflow^19^, to retrieve module specific expressions (AUCell scores) for each cell. We found trait (diagnosis, plaque stage, tangle stage) associated modules by fitting a linear regression model to predict the trait based on the module score, while regressing out the effects of sex. For each trait, multiple testing was controlled at FDR < 0.05 with the Benjamini-Hochberg procedure. The subnetwork for vesicle-mediated transport in module 2 was visualized using Cytoscape^68^. We determined marker genes using the Allen Brain Atlas whole brain Smartseq2 data^69^.

### Batch correction comparison

We compared top batch correction methods from Seurat, Harmony, and fastMNN with SCING module embeddings. To evaluate each method, we determined the average proportion of cells with the same group assignment (sample, batch, diagnosis, tangle stage, plaque stage, and sex), using 20 PCs and a variable number of neighbors (0.25, 0.5, 1, 2, 4, 8, and 16 percent of the dataset) (Equation 1). We determined the ability of each approach to remove batch and sample specific differences while retaining biologically relevant differences (diagnosis, tangle stage, plaque stage, and sex) by removing the batch and sample differences with an F1-score^43^ (Equation 2).

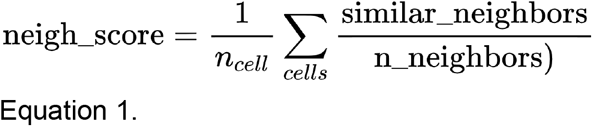

*neigh_score:* neighborhood score used to find the average fraction of neighbors of the same type (i.e. batch).
*similar_neighbors:* number of neighbors of a given cell that have the same identity (i.e. batch)
*n_neighbors:* number of total neighbors checked
*n_cell_*: total number of single cells

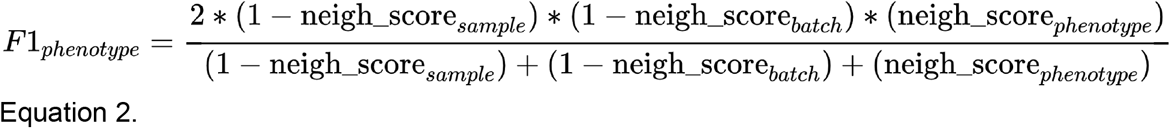

*neigh_score:* neighborhood score computed in Equation 1 for a given identity

### Application of SCING to Visium spatial transcriptomics data for mouse AD and WT brain

To determine the applicability and interpretability of SCING to spatial transcriptomics data, we applied SCING to mouse whole brain AD and WT data^28^. Since the network was built on the whole brain rather than a single cell type, we expect more variance amongst networks from subsamples, therefore we built 1,000 GRNs to be merged into the final network. We partitioned the genes with the Leiden graph partitioning algorithm into 33 modules. Using AUCell from SCENIC^19^, we obtained module specific expression for each spot. We determined regional specificity between pairs of larger regions (cortex, hippocampus, brainstem) through t-tests and overall variance for the smaller subregions through ANOVA. We determine differential module expression between AD and WT through t-tests, and correlation with age or plaque with Pearson correlation. Finally, the module 9, 10, 25, and 30 subnetworks were visualized using Cytoscape^68^. We determined marker genes using the Allen Brain Atlas whole brain Smartseq2 data^69^.

## Supplementary Figures

**Supplemental Figure 1.**
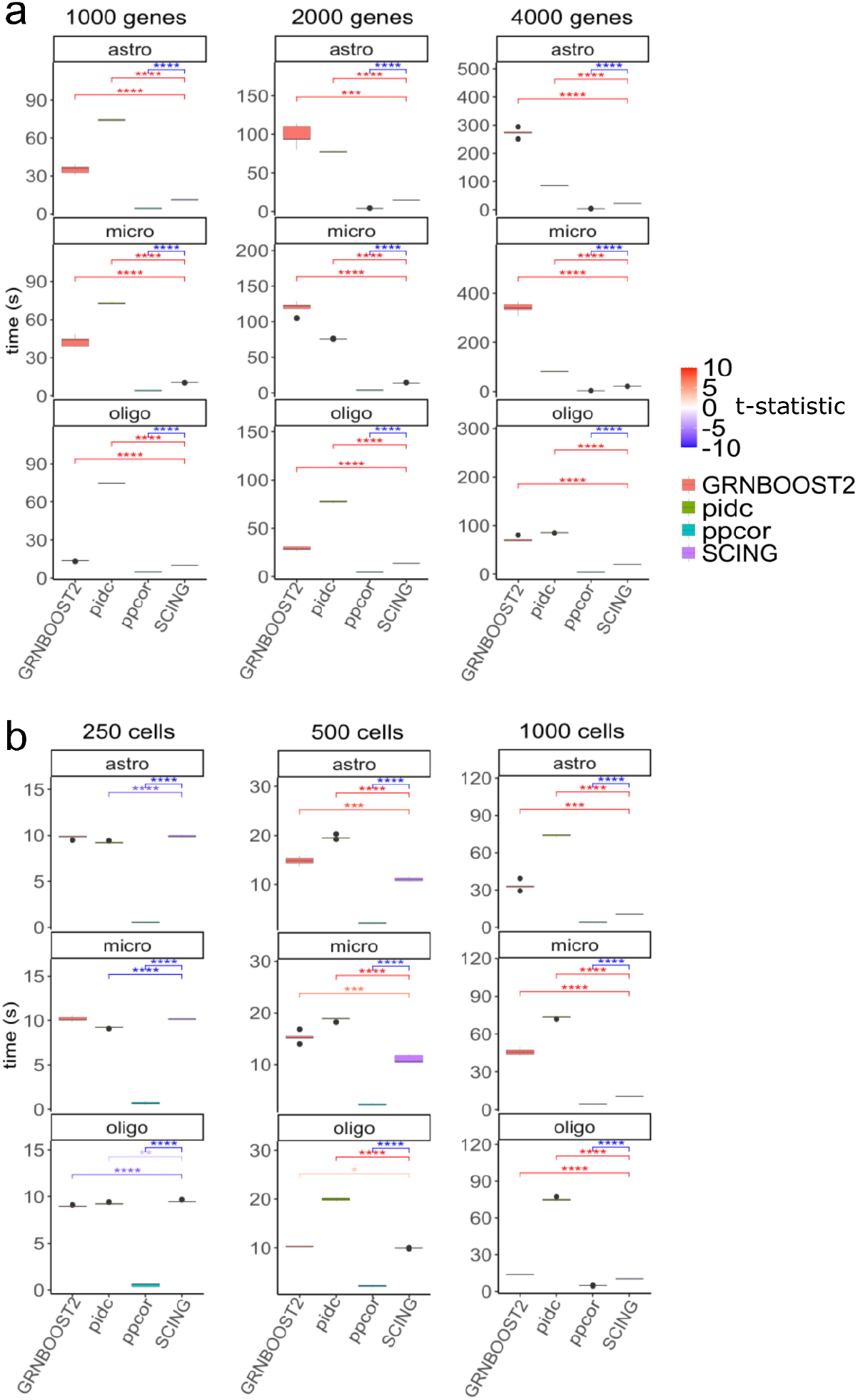
Run time comparison of methods for building a single GRN. SCING is faster than GRNBOOST2 and PIDC as the number of genes increases (a). The test increasing the number of genes was performed on 1000 cells. SCING is faster than GRNBOOST2 and PIDC as the number of cells increases (b). The test increasing the number of cells was performed on 1000 genes, ppcor is faster than all other methods. Red significance indicates that SCING is faster, while blue indicates SCING is slower. (*: p < 0.05, **: p < 0.01, ***: p < 0.001, ****: p < 0.0001)

**Supplemental Figure 2.**
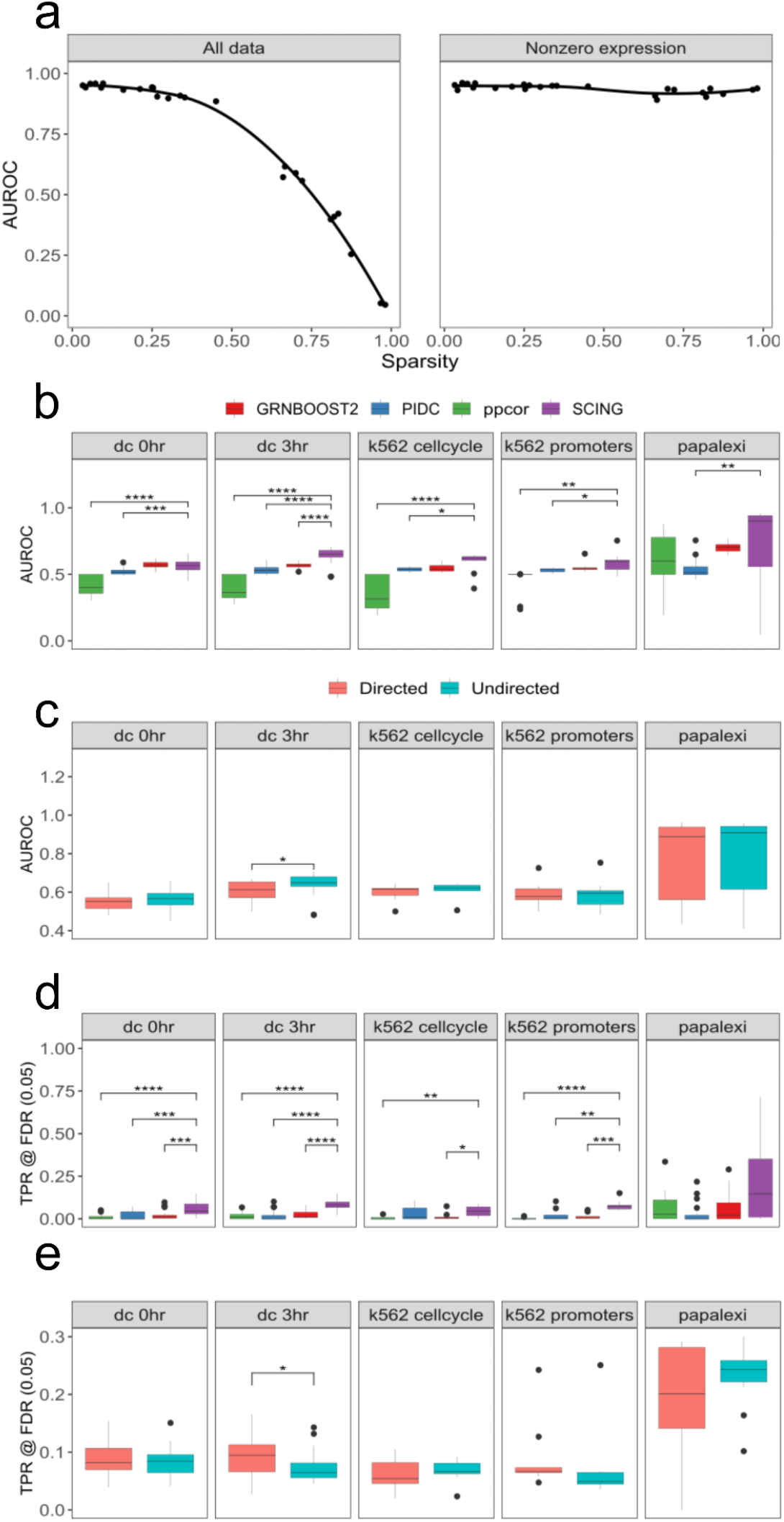
Predicted downstream affected genes of perturb-seq based perturbation in 5 datasets with GRNs built on all cells in each dataset. Performance of SCING in predicting downstream gene perturbation as a function of the fraction of zeros in each perturbation, before and after removing cells with zero expression of the gene of interest (a). Area under receiver operator characteristic (AUROC) curve for prediction of downstream perturbations using undirected GRNs (b). AUROC for prediction of downstream perturbation on directed GRNs for SCING (c). True positive rate (TPR) at a false discovery rate (FDR) of 0.05 for the prediction of downstream perturbations on undirected GRNs (d). TPR at FDR of 0.05 for the prediction of downstream perturbations on directed GRNs for SCING (e). (*: p<0.05, **: p<0.01,***: p < 0.001, ****: p< 0.0001)

**Supplemental Figure 3.**
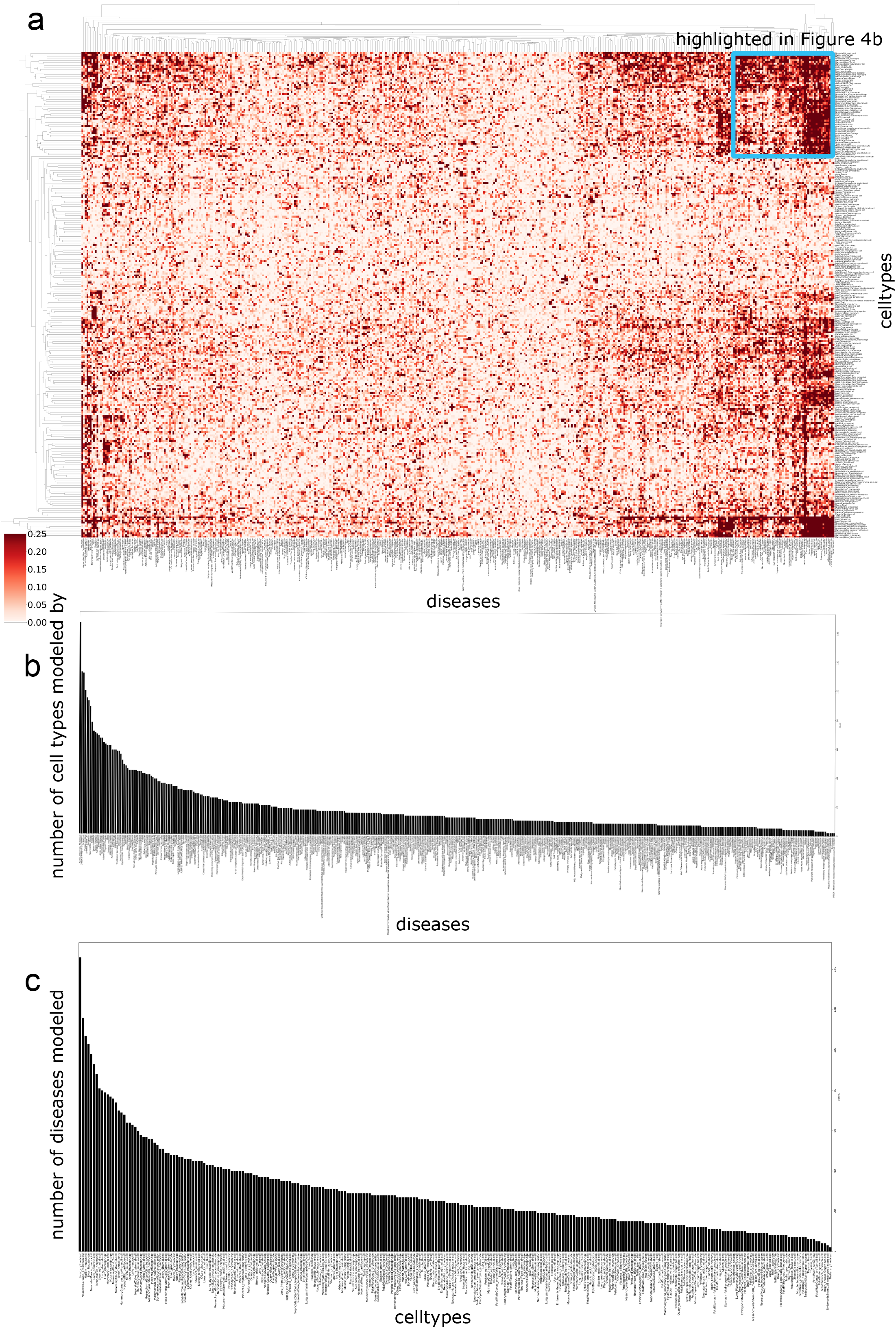
SCING GRNs’ modeling capabilities of DisGeNET diseases on entire MCA. Clustermap of all cell types in mouse cell atlas and SCING GRNs modeling of disease subnetworks across all diseases in DisGeNET (a). Number of cell types each disease subnetwork can be accurately (>0.1) modeled by (b). Number of disease subnetworks accurately modeled (> 0.1) by each cell type (c).

**Supplemental Figure 4.**
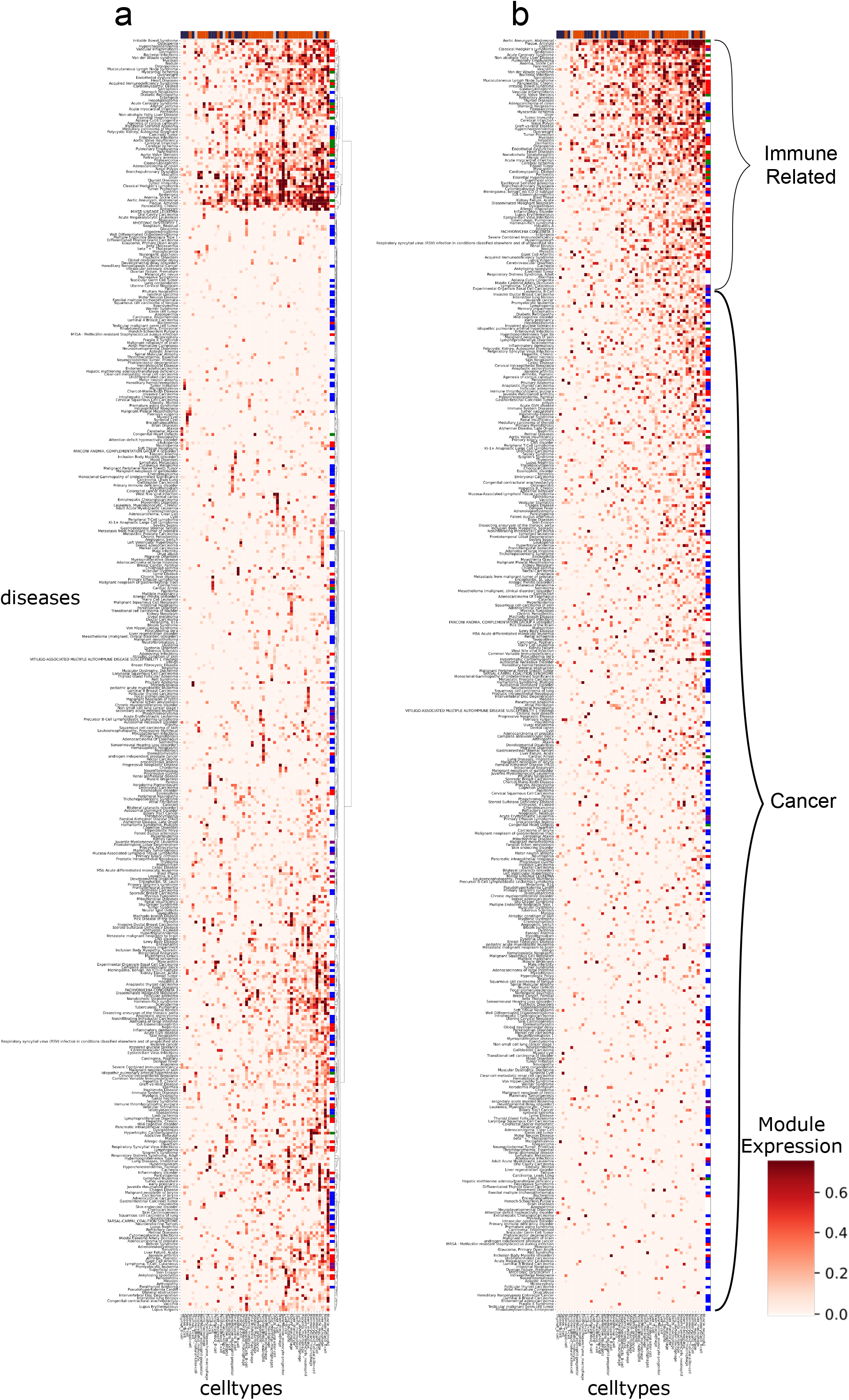
Clustermap of immune cell types and all diseases. Cell types are ordered by the number of diseases they accurately model, and diseases are either clustered with hierarchical clustering (a), or ordered by the number of cell types they are modeled by (b). Diseases are colored by disease category (immune related: red; cardiothoracic: green; cancer: blue; immune cell cancer: purple), and cell types are colored by innate (orange), and adaptive immune system (dark blue).

**Supplemental Figure 5.**
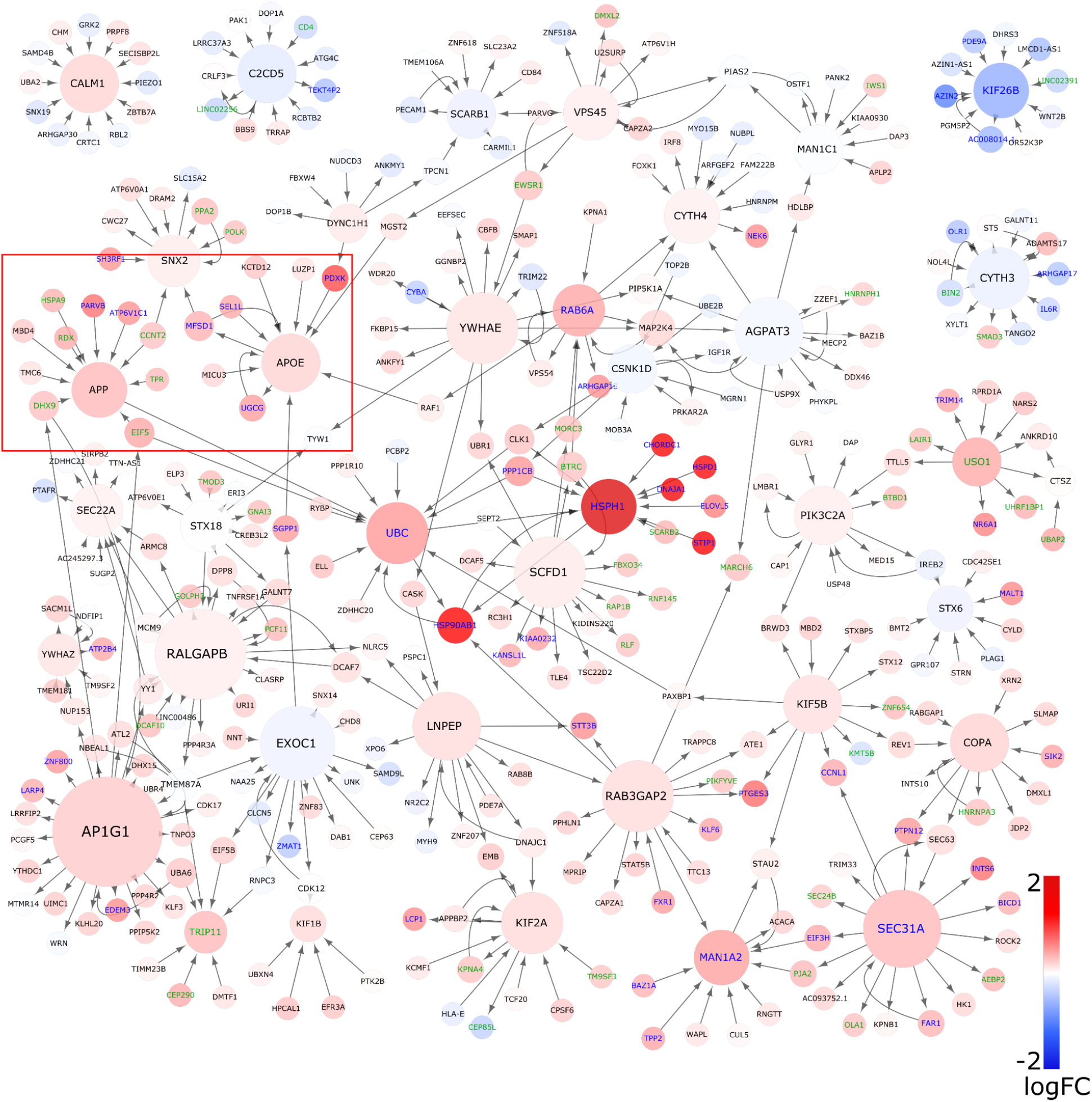
Subnetwork of the vesicle-mediated transport pathway from module 2 in the snRNAseq microglia GRN. Nodes are colored by their log-fold change (logFC) between AD and Control patients. APOE and APP subnetwork is highlighted (red box).

**Supplemental Figure 6.**
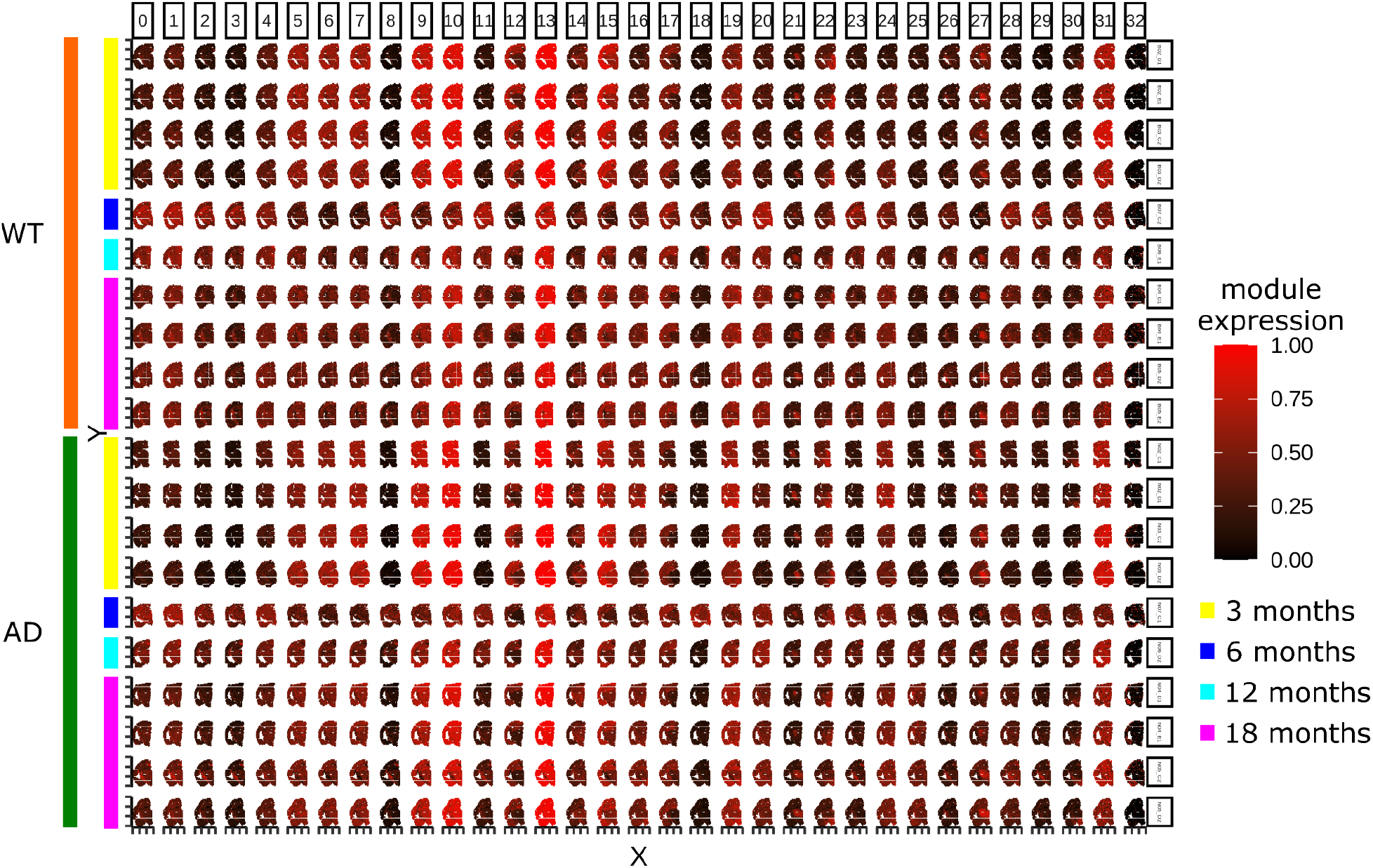
Expression of each module visualized in each brain. Mice are ordered by age (3, 6, 12, 18 months) and genotype (WT vs AD) mice.

**Supplemental Figure 7.**
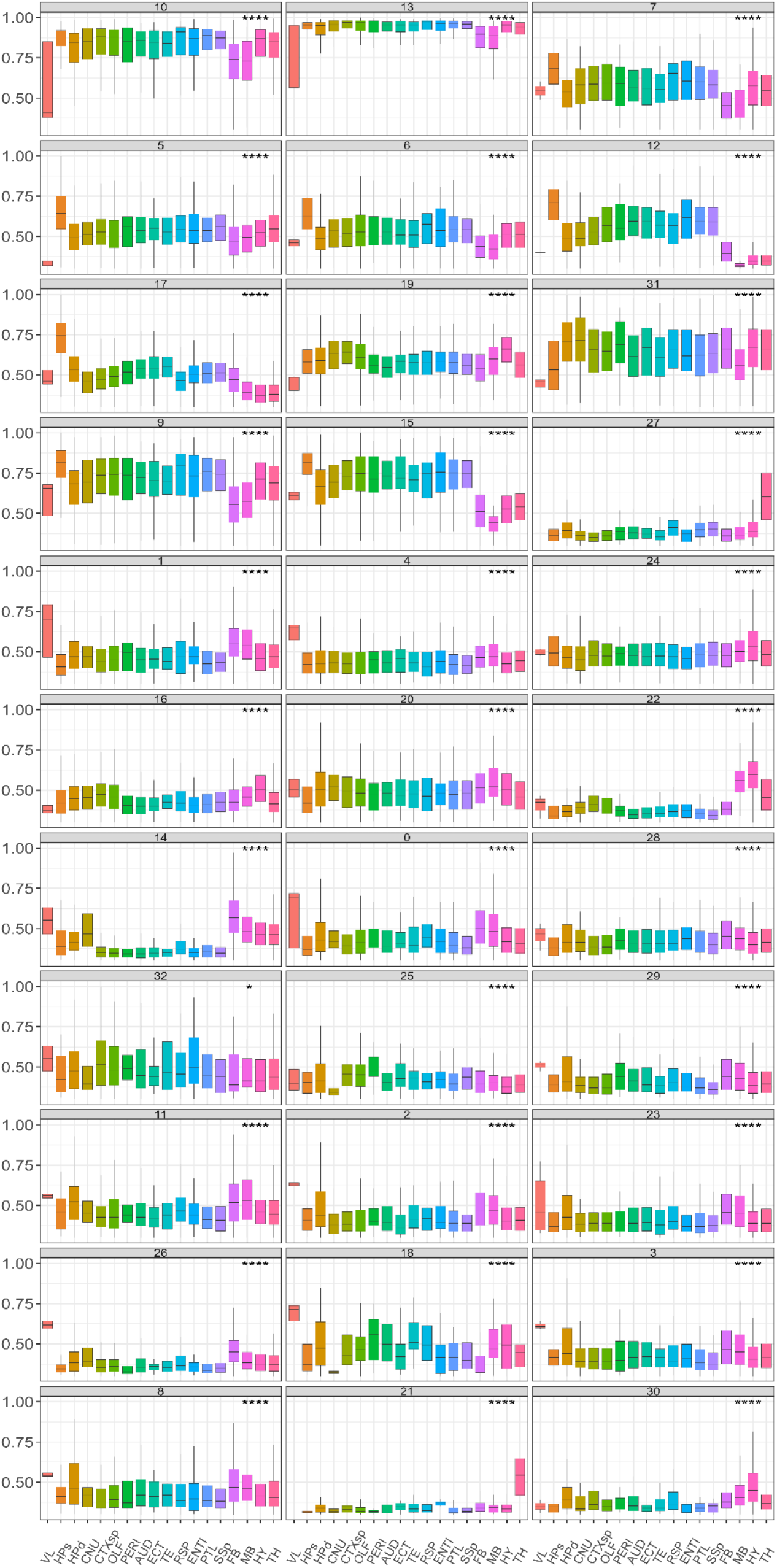
Distribution of module expression in each region of the brain. Significantly variable expression is determined by ANOVA (p<0.05).

**Supplemental Figure 8.**
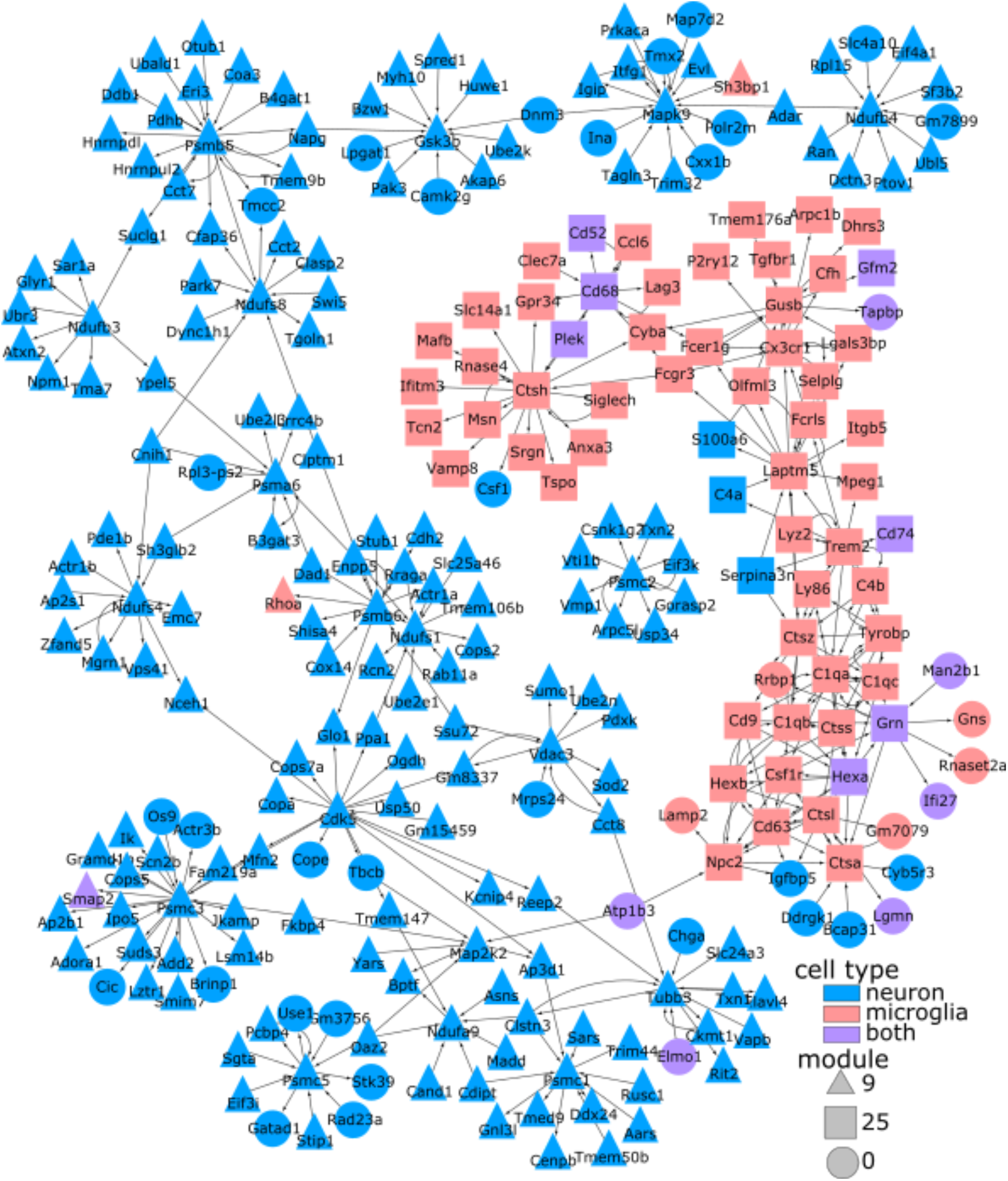
Subnetwork of module 9 (triangles) and 25 (squares), from the visium AD dataset. Marker genes were determined from the Allen Brain Atlas whole brain smart-seq dataset. Nodes are colored by marker gene status (microglia:red; neuronal:blue; both:purple). Cross cell type communication edges are between microglia and neuronal nodes. Module 9 is composed mostly of neuronal genes, while module 25 is composed mostly of mostly microglia genes.

**Supplemental Figure 9.**
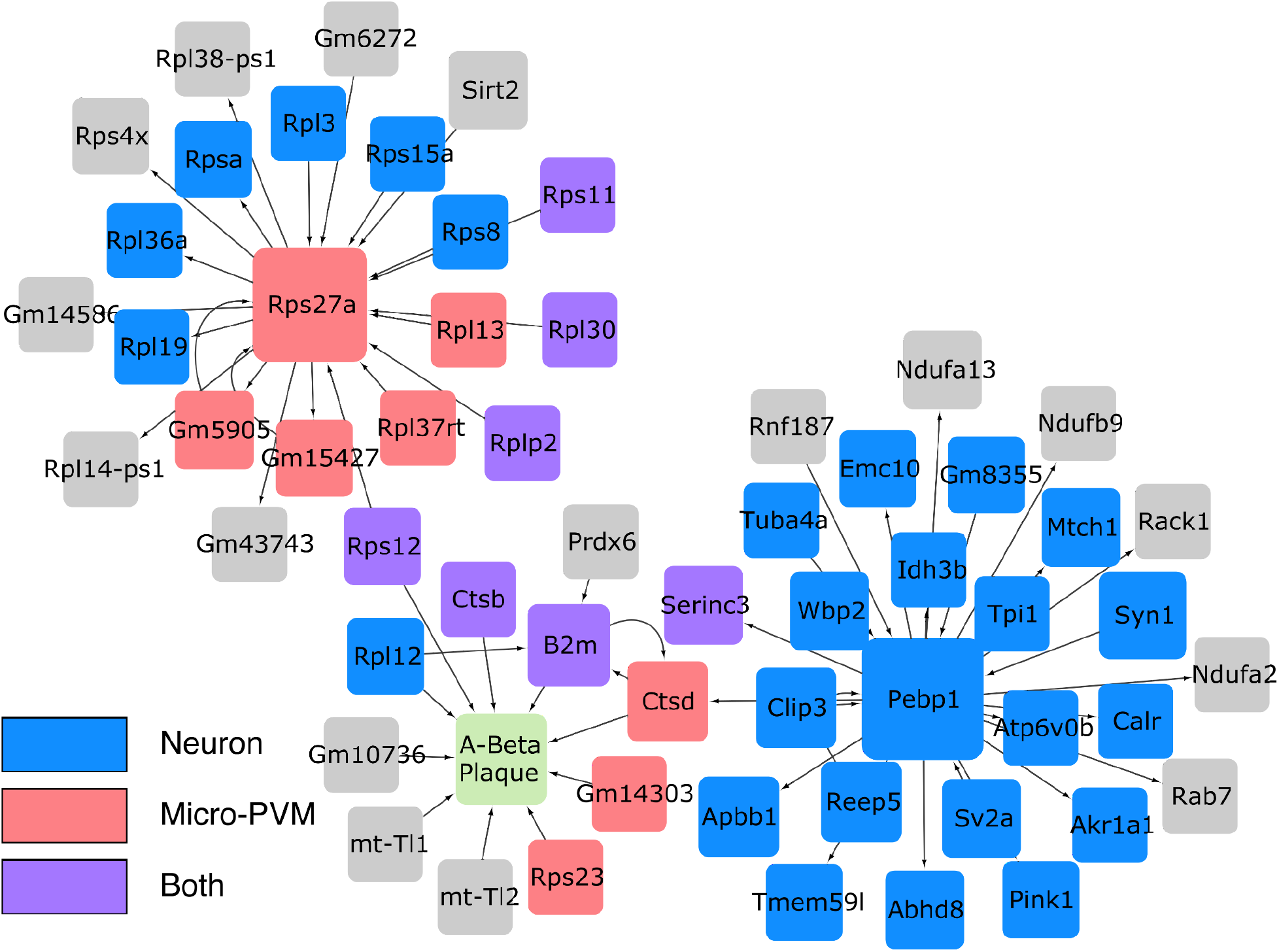
Subnetwork amyloid beta plaque in module 10 from the visium data. Nodes are colored by marker gene status (microglia:red; neuronal:blue; both:purple). Hypothesized cross cell type communication edges are between microglia and neuronal nodes. This contains the amyloid-beta plaque stain (colored green).

**Supplemental Figure 10.**
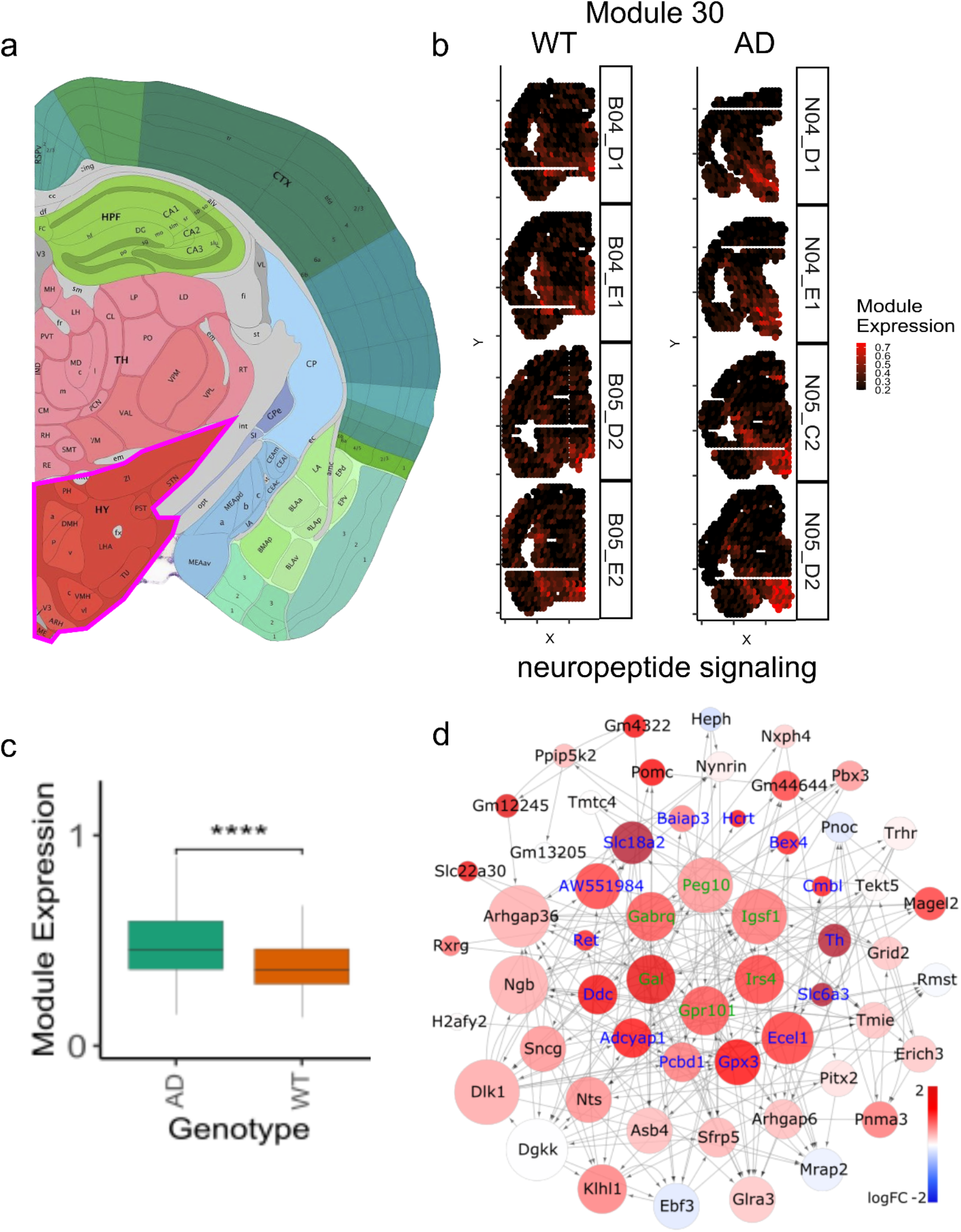
Localization and association of neuropeptide signaling in the hypothalamus. Diagram from the Allen Brain Atlas shows the location of the hypothalamus (outlined in pink) (a). Module 30 has localizes to the hypothalamus (b) and has higher expression in AD (c). The module 30 subnetwork (d) is enriched for neuropeptide signaling. Color of each node represents the logFC in AD vs WT mice. The color of the text represents significance (green: adjusted p-value < 0.05, blue: p-value < 0.01).

